# Resting-state functional connectivity predicts recovery from visually induced motion sickness

**DOI:** 10.1101/2020.06.30.180786

**Authors:** Jungo Miyazaki, Hiroki Yamamoto, Yoshikatsu Ichimura, Hiroyuki Yamashiro, Tomokazu Murase, Tetsuya Yamamoto, Masahiro Umeda, Toshihiro Higuchi

## Abstract

Movies depicting certain types of motion often provoke uncomfortable symptoms similar to motion sickness, termed visually induced motion sickness (VIMS). VIMS generally evolves slowly during the viewing of a motion stimulus and, when the stimulus is removed, the recovery proceeds over time. Recent human neuroimaging studies have provided new insights into the neural bases underlying the evolution of VIMS. In contrast, no study has investigated the neural bases underlying the recovery from VIMS. Study of the recovery process is critical for the development of a way to promote recovery and could provide further clues for understanding the mechanisms of VIMS. We thus investigated brain activity during the recovery from VIMS with functional connectivity (FC) magnetic resonance imaging. We found enhanced recovery-related FC patterns involving brain areas such as the insular, cingulate, and visual cortical regions, which have been suggested to play important roles in the emergence of VIMS. These regions also constituted large interactive networks. Furthermore, the increase in FC was correlated with the subjective awareness of recovery for the following 5 pairs of brain regions: insula–superior temporal gyrus, claustrum–left and right inferior parietal lobules, claustrum–superior temporal gyrus, and superior frontal gyrus–lentiform nucleus. Considering the previous findings on the functions of these regions and the present findings, it is suggested that the increase in FC may reflect brain processes such as enhanced interoceptive awareness to one’s own bodily state, a neuroplastic change in visual processing circuits, and/or the maintenance of visual spatial memory.

## Introduction

First-person perspective images, such as movies captured by action cameras or drones, have recently become popular. Such movies are so realistic that viewers can feel as if they themselves are moving inside the image space. However, the rich image experience is associated with a side effect, termed visually induced motion sickness (VIMS). VIMS is an unpleasant motion sickness (MS)-like symptom caused by movies depicting certain types of movement. The symptoms are classified into three types: (1) oculomotor (eyestrain, difficulty focusing, headache), (2) nausea (stomach awareness, increased salivation, burping), and (3) disorientation (dizziness, vertigo, drowsiness) (Shupak & Gordon, 2006; Kennedy *et al*., 2010). These symptoms can last for more than an hour (Kennedy *et al*., 1993a; Ujike *et al*., 2008).

VIMS generally emerges and evolves slowly during exposure to a motion stimulus, and, when the stimulus is removed, the individual slowly recovers from the MS symptoms over time. As for the emergency and evolutional phase of VIMS, several neuroimaging studies have revealed the underlying brain response (Napadow *et al*., 2013; Miyazaki *et al*., 2015; Farmer *et al*., 2015; Toschi *et al*., 2017). Napadow *et al*. (2013) measured the functional magnetic resonance imaging (fMRI) activity of female participants during the presentation of translating black/white stripes for a long enough period to induce VIMS, finding that nausea was associated with a broad network of brain areas including the insula, cingulate cortex, and limbic regions, which are known to process stress, emotion, and interoception. In addition, Miyazaki *et al*. (2015) measured fMRI visual cortical activities during the evolutional phase of VIMS, which was provoked by the presentation of actual images containing translational and rotational global motion components. The results showed desynchronized activities between the left and right inter-hemispheric middle temporal complex (MT+), which is a visual motion-sensitive area, when participants had MS. Taken together, it is suggested that various brain areas play key roles in the course of VIMS evolution.

Study of the processes of recovery from VIMS is critical for the development of a way to promote recovery from the unpleasant symptoms of VIMS and could also provide clues for further investigation of its mechanisms. The brain processes underlying the recovery phase of VIMS are not yet clear but previous studies on unpleasant events similar to MS, such as fatigue or stress, may provide some hints. For example, Peltier *et al*. (2005) reported a fatigue-related reduction in functional connectivity that was defined as a temporal correlation of brain activities. They measured fMRI time-series in the rest phase after participants completed a muscle fatigue task and found decreased functional connectivity between the motor cortices of the left and right hemispheres. As another example, van Marle *et al*. (2010) measured resting-state fMRI activity soon after female participants experienced psychological stress induced by viewing a strongly aversive movie. They reported increased functional coupling of the amygdala with the dorsal anterior cingulate cortex, anterior insula, and dorsorostral pontine region, indicating the extended state of hypervigilance that promotes sustained salience and mnemonic processing. These examples suggest that, during recovery, brain reorganisation occurs in functional brain networks related to the negative effects.

We therefore conjectured that such brain reorganisation would also occur during the recovery from VIMS. To test this prediction, we investigated resting-state fMRI activities during the recovery phase of VIMS. Specifically, we predicted that the functional connectivity of brain regions such as the insula, cingulate, and/or visual cortical regions would be selectively changed because these regions were speculated to play important roles in the emergency and evolutional phase of VIMS.

## Methods

### Participants

Participants were 14 volunteers (12 men and 2 women, including 4 authors; mean age, 34.9 years; range, 25–48 years) with normal or corrected-to-normal vision. All participants had no history of neurologic or psychiatric disorder and provided written informed consent. All experimental procedures were approved by the Human Studies Committee of the Graduate School of Human and Environmental Studies at Kyoto University and the Department of Neurosurgery at Meiji University of Oriental Medicine.

According to the previous study (Miyazaki *et al*., 2015), the participants were classified into two groups: the VIMS group (n = 8, all men, including 2 authors; mean age ± SD, 32.3 ± 7.2 years) who experienced VIMS in the experimental session; and the healthy group (n = 6, 2 women and 4 men including 2 authors; mean age ± SD, 33.3 ± 7.2 years) who did not get VIMS. Details on this grouping criterion are provided in Miyazaki *et al*. (2015). Two in the VIMS group could not recover during the experimental session and were thus excluded from this analysis aimed at investigating the recovery phase from VIMS. Consequently, six VIMS participants (n = 6, all men including 2 authors; mean age ± SD, 34.7 ± 8.0 years) were included in the analysis.

### Visual stimuli

In the experiment, two types of visual stimuli—a global motion stimulus and a local one—were presented. The global motion stimulus was a 6-minute-long actual movie containing first-person-perspective, global visual motion. This stimulus content could induce viewers’ VIMS. In contrast, the local motion stimulus was a 6-minute-long actual movie consisting of 8 by 8 patches (total, 64), which reduced the global motion stimulus to one-eighth vertically and horizontally. This stimulus was a control and did not induce viewers’ VIMS because the stimulus contained no global motion component that could trigger VIMS. The stimulus was based on Wall & Smith (2008).

These stimuli were projected onto a screen located over the participant’s forehead using a DLP projector (LVP-HC3800; Mitsubishi, Japan). The spatial resolution was 1080 × 720 pixels, and the refresh rate was 60 Hz. The participant was supine and viewed the stimuli through a planar mirror located 25 cm from the eyes. See Miyazaki *et al*. (2015) for details.

### Experimental procedure

The experimental procedure is shown in Figure 1. Before and after the presentation of the two aforementioned stimuli, three rest phases (Rest-1, Rest-2, and Rest-3) were arranged. Each rest phase was 5 minutes long and comprised presentation of a grey background with a fixation point at the centre. During the rest phases, participants were asked to relax with their eyes open. After the first rest phase (Rest-1), participants viewed the local motion stimulus; none got VIMS. Next, the participants experienced the second rest phase (Rest-2) and then viewed the global motion stimulus; 8 of the 14 participants got VIMS, as described above. The third rest phase (Rest-3) followed. The order of the stimulus presentations (the local motion stimulus followed by the global one) was not balanced to avoid a carryover effect from the global to the local motion period because our preliminary experiment had indicated that, when the global stimulus precedes the local one, the VIMS symptoms can be prolonged and overlap the subsequent local stimulus presentation.

**Figure 1.**
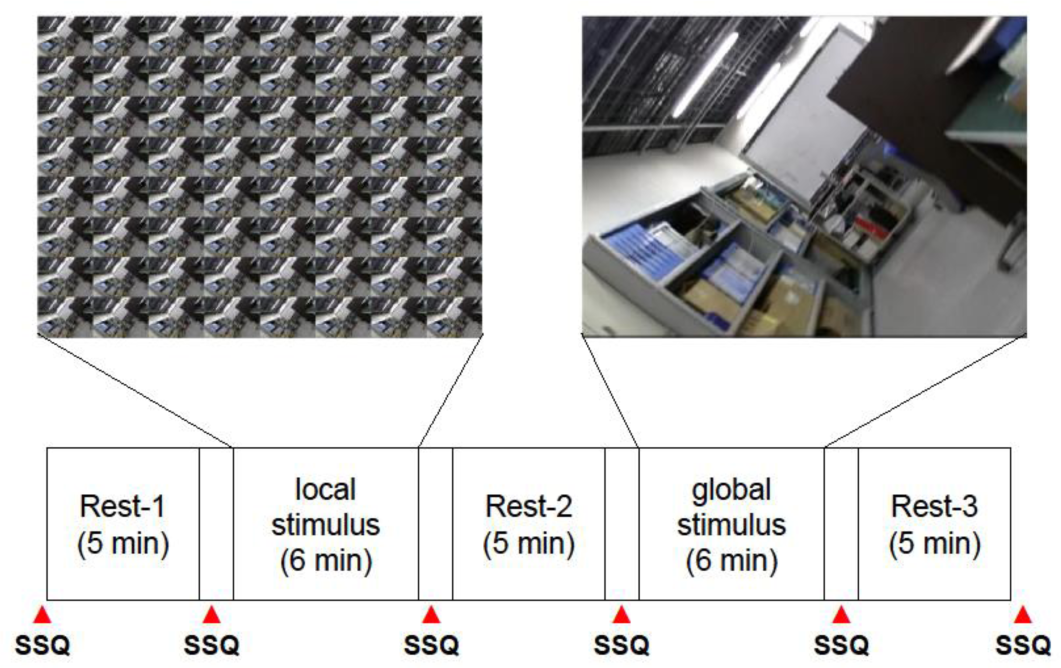
Schematic diagram showing the time course of the experimental session. After the first rest phase (Rest-1), the local motion stimulus was presented as a control. After the second rest phase (Rest-2), the global motion stimulus, which can induce visually induced motion sickness (VIMS), was presented. Finally, the third rest phase (Rest-3) comprised the recovery from VIMS.

To rate the degree of VIMS symptoms, participants answered the Simulator Sickness Questionnaire (SSQ) (Kennedy *et al*., 1993b) 6 times in total, that is, before and after each stimulus presentation and each rest phase. The SSQ was developed as a metric of simulator sickness, and this metric consists of 16 questions on symptoms such as headache and nausea, based on four grades. The SSQ has generally also been used as a metric for VIMS. The question items of SSQ were presented to participants through images, and the questions were answered by use of a response pad with four buttons (HH-1×4L; Current Design Inc., Philadelphia, PA). A decrease in the SSQ scores between before and after the rest phases, which would reflect recovery from VIMS, was analysed because the aim of the present study was to investigate the recovery phase of VIMS at rest.

### MRI data acquisition

Functional MR images were acquired using a clinical 3T-MR scanner (Trio TIM; Siemens, Germany) and a 20-channel phased-array head coil with a T2^*^-weighted gradient-echo echo planer sequence (repetition time [TR] = 2000 ms; echo time [TE] = 30 ms; flip angle [FA] = 90°; voxel size [VS] = 3 × 3 × 4 mm^3^; field of view [FOV] = 192 × 192 mm^2^; 37 slices; axial orientation parallel to the AC-PC plane; interleaved acquisition). The functional images were continuously scanned from 10 seconds before Rest-1 to the end of Rest-3, taking about 35 minutes. For anatomical registration, brain structural images were acquired using a 3D magnetization-prepared rapid gradient-echo T1-weighted sequence with the following parameters: TR = 1800 ms, TE = 3.03 ms, inversion time = 650 ms, FA = 9°, VS = 0.8 × 0.8 × 0.8 mm^3^, FOV = 205 × 205 mm^2^, 176 slices, axial orientation parallel to the AC-PC plane, interleaved acquisition.

### Behavioural data analyses

#### Visually induced motion sickness

Participants self-reported whether, when viewing the global motion stimulus, they experienced VIMS or not after the experimental session and, based on the self-report, were provisionally classified into the VIMS or healthy group. To objectively test the validity of this grouping criterion based on the self-reports, the scores of the SSQ, which is a metric for MS symptoms, were statistically tested to determine whether the scores of the VIMS group selectively increased after the global motion stimulus. Details on this analysis are provided in Miyazaki *et al*. (2015).

#### Recovery from visually induced motion sickness

Next, to verify that the VIMS group participants recovered during Rest-3, the differences in the SSQ scores between before and after each rest phase were computed, and the differences were statistically tested to determine whether the score of the VIMS group became significantly larger only for Rest-3, reflecting the recovery from VIMS, and whether the scores of the other rest phases did not change. This analysis was performed using R software (R Core Team, 2017) (see also Miyazaki *et al*., 2015). A linear mixed-effects model analysis was performed with the *lmer* function of the lme4 package (Bates *et al*., 2014). This model had a within-participants fixed effect PHASE (Rest-1 *vs*. Rest-2 *vs*. Rest-3), a between-participants fixed effect GROUP (VIMS *vs*. healthy participant groups), an interaction of PHASE and GROUP, and a random effect of each individual participant. This analysis was conducted separately for the four scales of the SSQ (Total Score, Nausea, Oculomotor, and Disorientation).

When the PHASE × GROUP interaction was statistically significant, for the *post hoc* analysis, the simple effect of PHASE was tested separately for the VIMS and healthy groups. Specifically, the differences in scores between the levels of PHASE (i.e., three ways of Rest-1 *vs*. Rest-2; Rest-2 *vs*. Rest-3; and Rest-1 *vs*. Rest-3) were tested by means of the functions *testInteractions* and *testFactors* of the phia package (De Rosario-Martinez, 2015) separately for each participant’s group. In contrast, when the interaction was not significant, the main effect of PHASE was reported instead. The significance level for all statistical analyses was set to 0.05, with correction by the Bonferroni method for multiple comparisons involving the four scales of the SSQ, the two participant groups, and the repetition of the *post hoc* tests.

### fMRI data analyses

#### Preprocessing

All fMRI data were preprocessed using AFNI software (Cox, 1996) as follows: (1) slice timing correction; (2) head motion correction; (3) extraction of 5-minute-long time-series signals (150 fMRI volumes) acquired during each rest phase; (4) transformation to the standard Talairach space for group analysis; (5) smoothing with the Gaussian kernel of an isotropic 9-mm FWHM; (6) band-pass filtering with a cutoff of 0.01–0.1 Hz; and (7) removal of fMRI volumes whose motion or its derivatives exceeded 0.2 mm (i.e., motion ‘scrubbing’) to reduce the effect of head movement on fMRI data, in addition to the removal of outlier volumes where the ratio of the number of outlier voxels exceeded 0.1.

#### Brain connectedness mapping

After the preprocessing, connectedness maps were made by use of the AFNI *3dTcorrMap* function. Connectedness is the average of the Pearson correlation coefficients between a specific voxel time-series and the other in the brain mask (Gotts *et al*., 2012). The Pearson correlation coefficient is a real number ranging from –1 to +1, and this value was converted into a Fisher’s Z value with the *3dTCorrMap* function for the sake of the following analyses. The Z values were compared between the VIMS and healthy groups within the Talairach space by using the AFNI *3dLME* function. For the mapping of the results of this comparison, the voxel-wise *P* value was set to 0.0001. The false-positive rate for the clustering was estimated by use of the *3dClustSim* function, in which the threshold α for the cluster size was set to 0.10; this relatively loose threshold was used because this was a first screening procedure to restrict candidates. The conventional 0.05 threshold was used in the second screening based on functional connectivity described below. Based on this mapping, the regions whose connectedness changed significantly during the recovery phase of VIMS were detected. Specifically, the regions were determined according to the following criteria: the connectedness was changed only for the VIMS group selectively during Rest-3 when the VIMS participants got VIMS, whereas that of the healthy group did not change for all rest phases (interaction contrast calculated using the *3dLME* function: absolute intergroup difference in Z_Rest-3_ – 0.5 × Z_Rest-1_ – 0.5 × Z_Rest-2_: the VIMS group – the healthy group > 3.891, which corresponds to *P* < 0.0001).

Through these steps, 12 brain regions were identified with significant changes in connectedness during recovery from VIMS (described below). In each region, the mean time-series averaged across voxels within the region was calculated and used as a seed for the functional connectivity analysis in the following section.

#### ROI-based functional connectivity analysis

As stated above, the mean time-series averaged across voxels within the brain regions showing significant changes in connectedness during recovery from VIMS were used as seeds to examine functional connectivity through the entire brain. Then, using the same method used in the analysis of connectedness, the functional connectivity intensity was compared between the VIMS and healthy groups. After adjustment to the false-positive rate for clustering, the result was mapped in Talairach space (voxel-wise *P* value = 0.0001, α = 0.05), through which the brain regions with changes in functional connectivity during recovery from VIMS that fulfilled the following criteria were identified: (i) functional connectivity in Rest-3 following the onset of VIMS changed only in the VIMS group (interaction contrast calculated using the *3dLME* function: absolute intergroup difference in Z_Rest-3_ – 0.5 × Z_Rest-1_ – 0.5 × Z_Rest-2_: the VIMS group–the healthy group > 3.891, which corresponds to *P* < 0.0001) and (ii) neither group had a change in functional connectivity during other rest phases (the VIMS group, |Z_Rest-2_ – Z_Rest-1_| < 1.96, which corresponds to *P* > 0.05; the healthy group, |Z_Rest-3_ – 0.5 × Z_Rest-1_ – 0.5 × Z_Rest-2_| < 1.96; |Z_Rest-2_ – Z_Rest-1_| < 1.96).

This functional connectivity analysis extracted 19 pairs of brain regions, and changes in each functional connectivity were analysed using R software (R Core Team, 2017) and a linear mixed-effects model (the same model used in the analysis of the SSQ scores) consisting of a within-participants fixed effect (PHASE; Rest-1 *vs*. Rest-2 *vs*. Rest-3), a between-participants fixed effect (GROUP; VIMS *vs*. the healthy group), an interaction of PHASE and GROUP, and a random effect of each individual participant.

Since the PHASE × GROUP interactions for all region pairs were statistically significant, a *post hoc* test was performed to analyse the simple main effect of PHASE in the VIMS and healthy groups separately. Specifically, the differences in connectivity between Rest-1 and Rest-2, Rest-2 and Rest-3, and Rest-1 and Rest-3 were tested in each group. The Bonferroni method was used to correct multiple comparisons for functional connectivities in the 19 pairs of brain regions, the two participant groups, and repeated *post hoc* tests of the three PHASE combinations.

#### Correlation analysis

Correlation analysis was performed to clarify whether a change in SSQ Total Score (i.e., subjective awareness of recovery) could be predicted from the change in functional connectivity. Because exceptionally large changes in SSQ Total Scores were observed in some participants, the Spearman’s rank correlation coefficient, a nonparametric statistic that is seldom affected by such variation, was used to analyse the 19 pairs of brain regions separately. Because of the multiple comparisons, *P* values were subjected to Bonferroni correction.

#### Global network analyses

To analyse the functional network of the brain regions showing recovery-related increases in connectedness, a multivariate analysis was performed. First, for each participant and for each rest phase, a partial correlation coefficient of the fMRI time-series was computed for each pair of the 12 brain regions, resulting in a 12 × 12 matrix. The partial correlation matrix was then converted into a distance matrix, where the distance was defined by 1 minus the absolute partial correlation coefficient. Second, for each rest phase, within-group averaged distance matrices were computed for VIMS and healthy groups. The distance matrices of the first phase (Rest-1) and second phase (Rest-2) were further averaged as a control for the recovery phase (Rest-3). Third, the columns and rows of the distance matrices were reordered to place similar brain regions closer, using the *seriation* function with the ‘HC’ option in the *seriation* package (Hahsler *et al*. 2008) of R software. Finally, dendrograms, which are hierarchical clustering trees, were derived from the distance matrices using the *pvclust* package (Suzuki & Shimodaira 2019) of R software, which provided *P* values of each cluster through multiscale bootstrap resampling. Independently of the above analyses, statistically independent brain networks were extracted by a dictionary learning framework. The procedure and the results are detailed in the Supplementary Materials.

## Results

### Behavioural results

Details of the results for VIMS symptoms induced in the experiment are provided in Miyazaki *et al*. (2015). In brief, 8 of the 14 participants experienced VIMS due to the global motion stimulus, and the remaining 6 did not (Figs. 3 and 4 of Miyazaki *et al*. [2015]). These results were confirmed by statistical analysis of the SSQ scores (Table 1 of Miyazaki *et al*. [2015]). Participants were divided into VIMS (n = 8) and healthy (n = 6) groups based on the presence and absence of VIMS, respectively. However, 2 of the 8 participants in the VIMS group did not recover from the VIMS even after the experimental session, and this was reflected in their SSQ scores, which did not decrease even after Rest-3. Because the aim of this study was to analyse the recovery phase of VIMS, these two participants were excluded from the subsequent analyses.

**Table 1.**
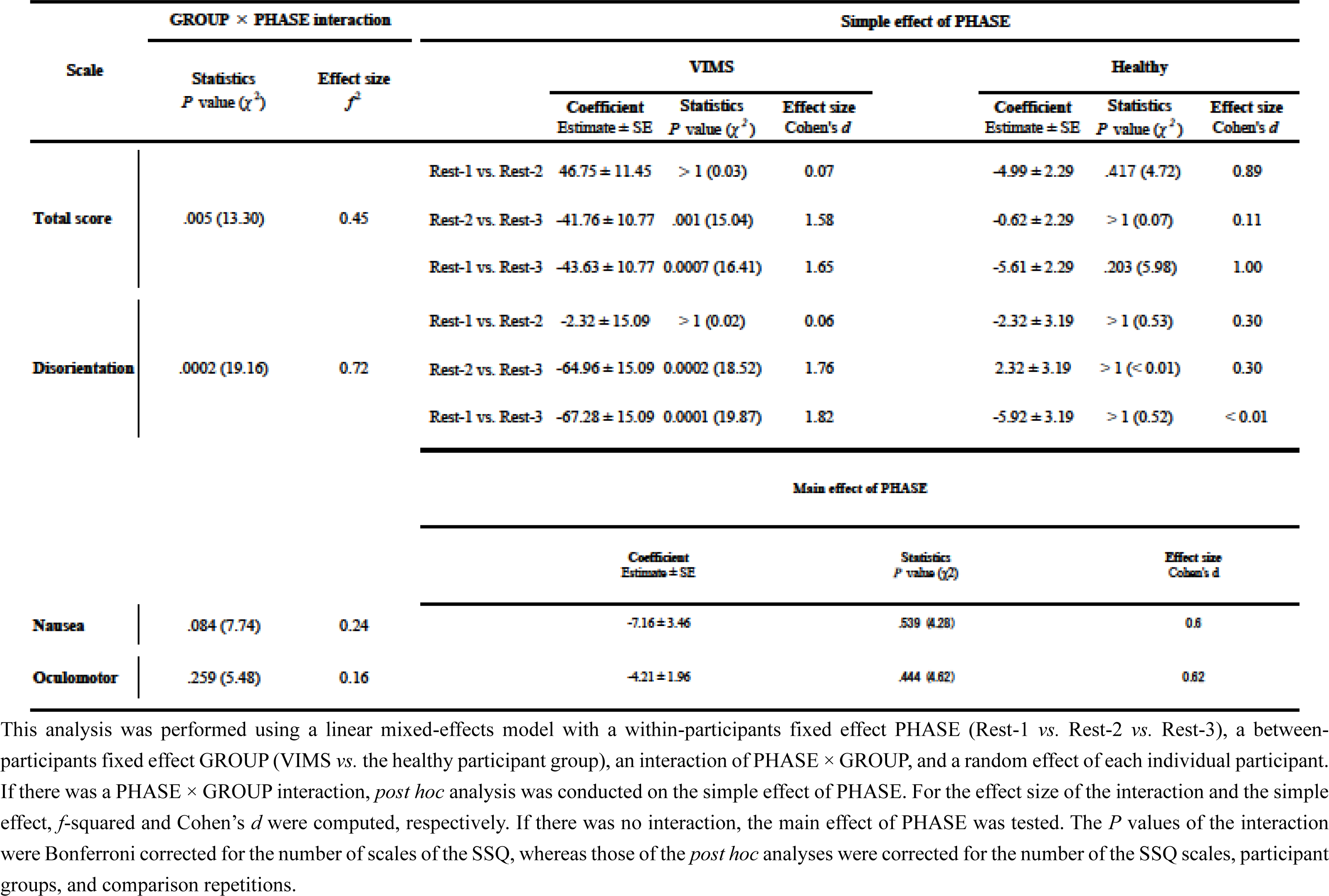
Statistical analysis of the degree of recovery in the subjective score of the Simulator Sickness Questionnaire (SSQ) (decrease in the SSQ between the post- and pre-rest phases).

**Figure 2.**
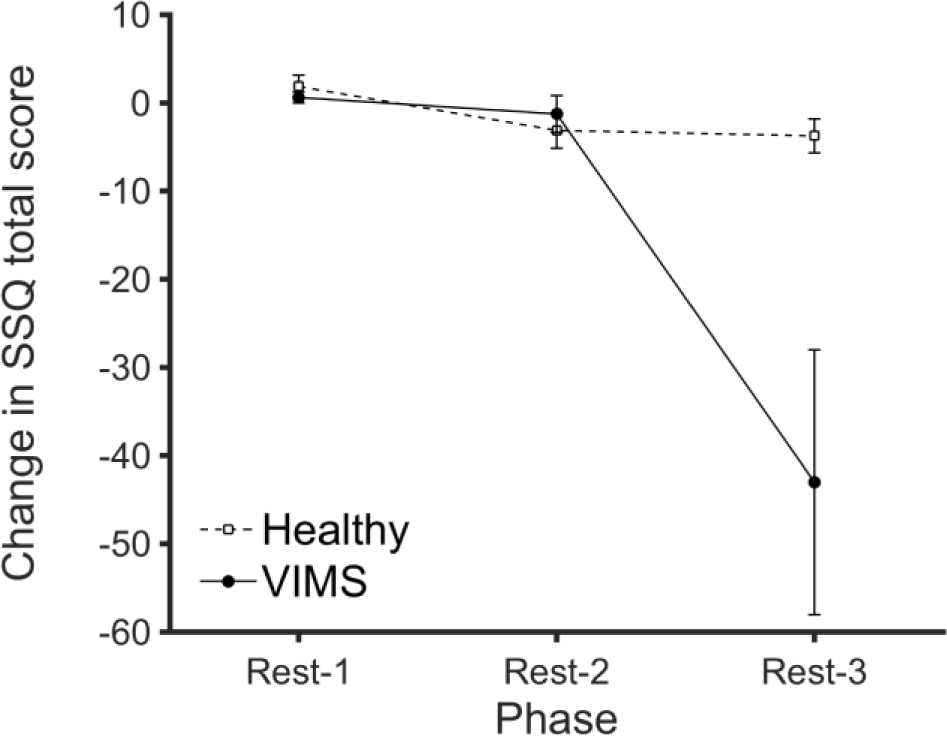
Time course of the change in the Total Score (TS) of the Simulator Sickness Questionnaire (SSQ). The vertical axis shows the change in the TS (post-score minus pre-score) of the SSQ. The horizontal axis shows the experimental phase (Rest-1, Rest-2, and Rest-3). The visually induced motion sickness (VIMS) group’s participants experienced motion sickness soon before the third rest phase (Rest-3), and recovery from symptoms occurred during Rest-3. Accordingly, the score of the VIMS group (solid line) significantly decreased before and after Rest-3, whereas the score of the healthy group participants (broken line), who did not experience VIMS, did not change as much.

**Figure 3.**
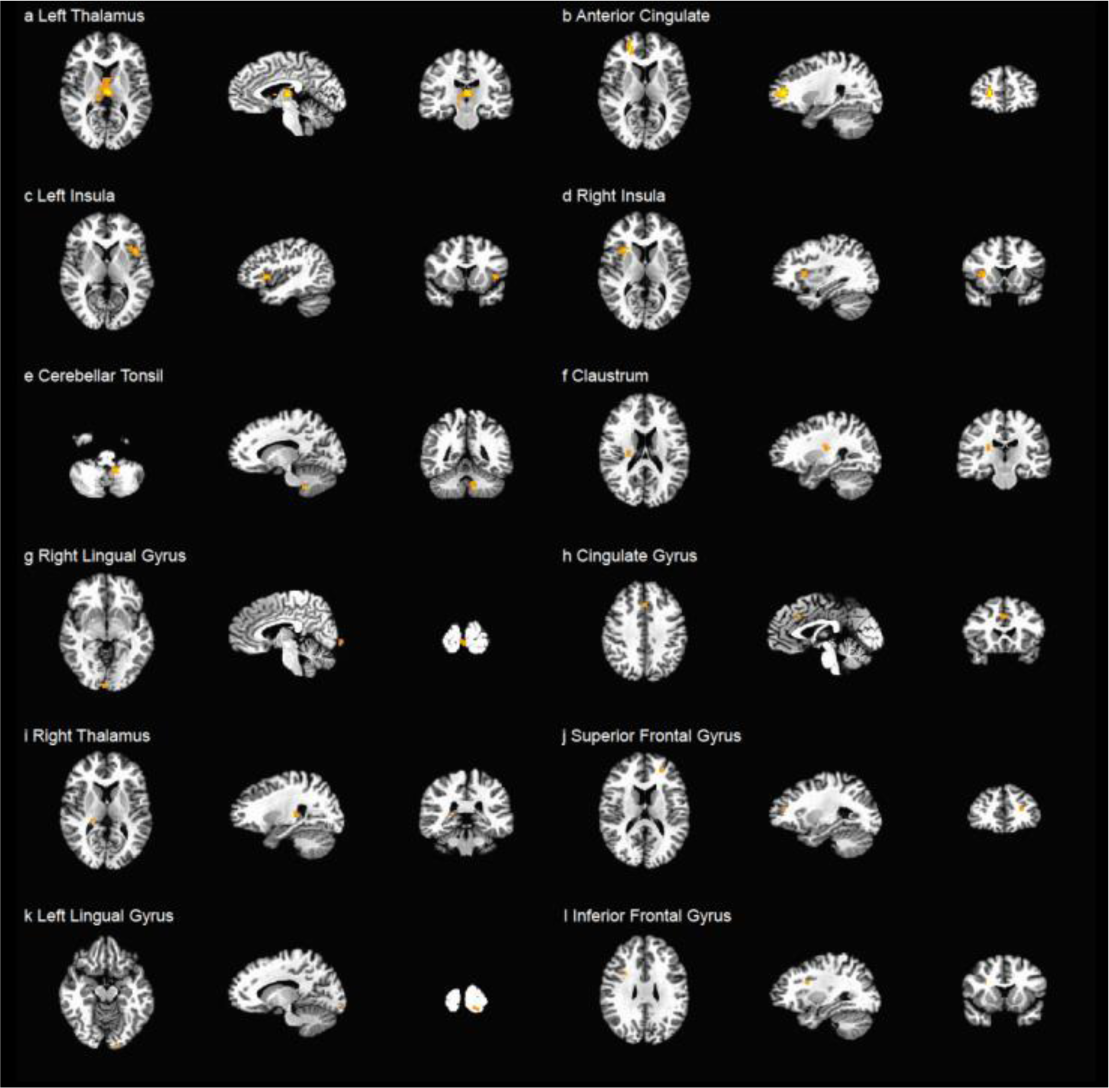
Twelve brain regions whose connectedness significantly increased during the recovery phase from visually induced motion sickness (VIMS).

**Figure 4.**
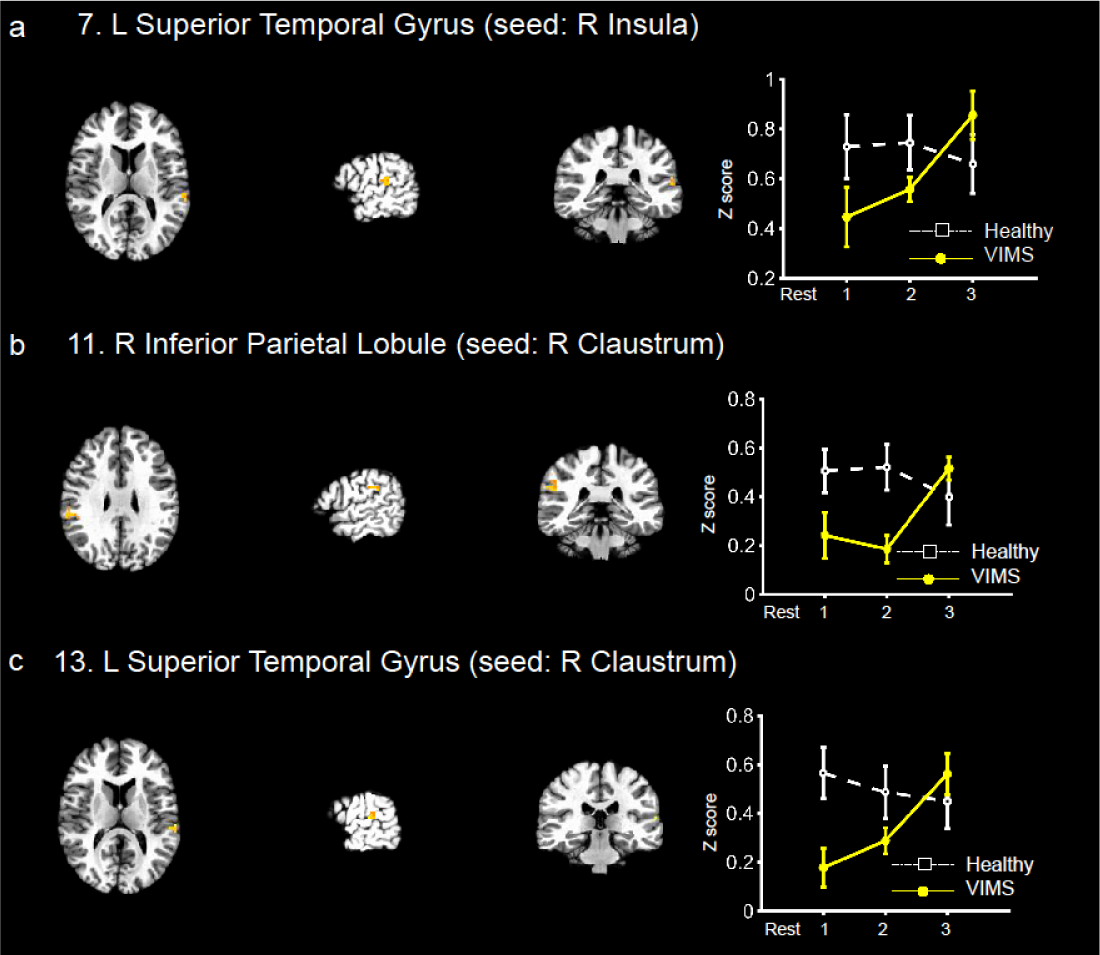
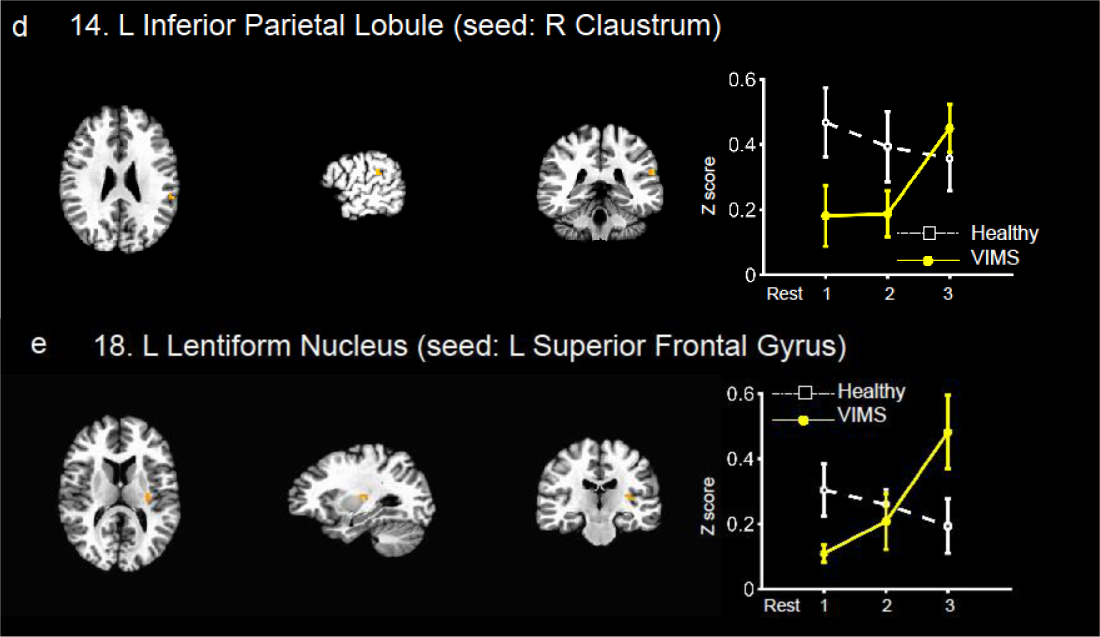
Brain regions whose functional connectivity with the seed regions increased in the recovery phase from visually induced motion sickness (VIMS). Each column shows, from left to right, axial, sagittal, and coronal brain images, and the change in the functional connectivity (Z scores averaged within the participants’ groups) of each experimental phase, separately for VIMS and healthy groups, respectively (error bars are standard errors). Each panel is the result for the following brain regions (the brain region in brackets is the corresponding seed): a–left superior temporal gyrus (right insula), b–right inferior parietal lobule (right claustrum), c–left superior temporal gyrus (right claustrum), d– left inferior parietal lobule (right claustrum); and e–left lentiform nucleus (left superior frontal gyrus). For this mapping, we used the following thresholds: voxel-wise threshold *P* < 0.0001, cluster level threshold *α* < 0.05. The numbering of brain region pairs corresponds to that in Tables 3, 4, and 5.

To verify that the participants in the VIMS group had recovered from VIMS during Rest-3, a linear mixed-effects model analysis of SSQ scores was performed before and after each rest phase (Table 1 and Fig. 2). SSQ Total Scores decreased during Rest-3 only in the VIMS group. The interaction of PHASE and GROUP was statistically significant (χ^2^(2) = 13.30, *P* = 0.005, Bonferroni corrected). A *post hoc* test of the simple main effect of PHASE showed that SSQ scores decreased significantly during Rest-3 only in the VIMS group (the VIMS group: Rest-1 *vs*. Rest-2: χ^2^(2) = 0.03, *P* > 1; Rest-2 *vs*. Rest-3: χ^2^(2) = 15.04, *P* = 0.001; Rest-1 *vs*. Rest-3: χ^2^(2) = 16.41, *P* = 0.0007; the healthy group: Rest-1 *vs*. Rest-2: χ^2^(2) = 4.72, *P* = 0.417; Rest-2 *vs*. Rest-3: χ^2^(2) = 0.07, *P* > 1; Rest-1 *vs*. Rest-3: χ^2^(2) = 5.98, *P* = 0.203, Bonferroni corrected), verifying that the participants in the VIMS group recovered from the VIMS during Rest-3.

**Table 2.**
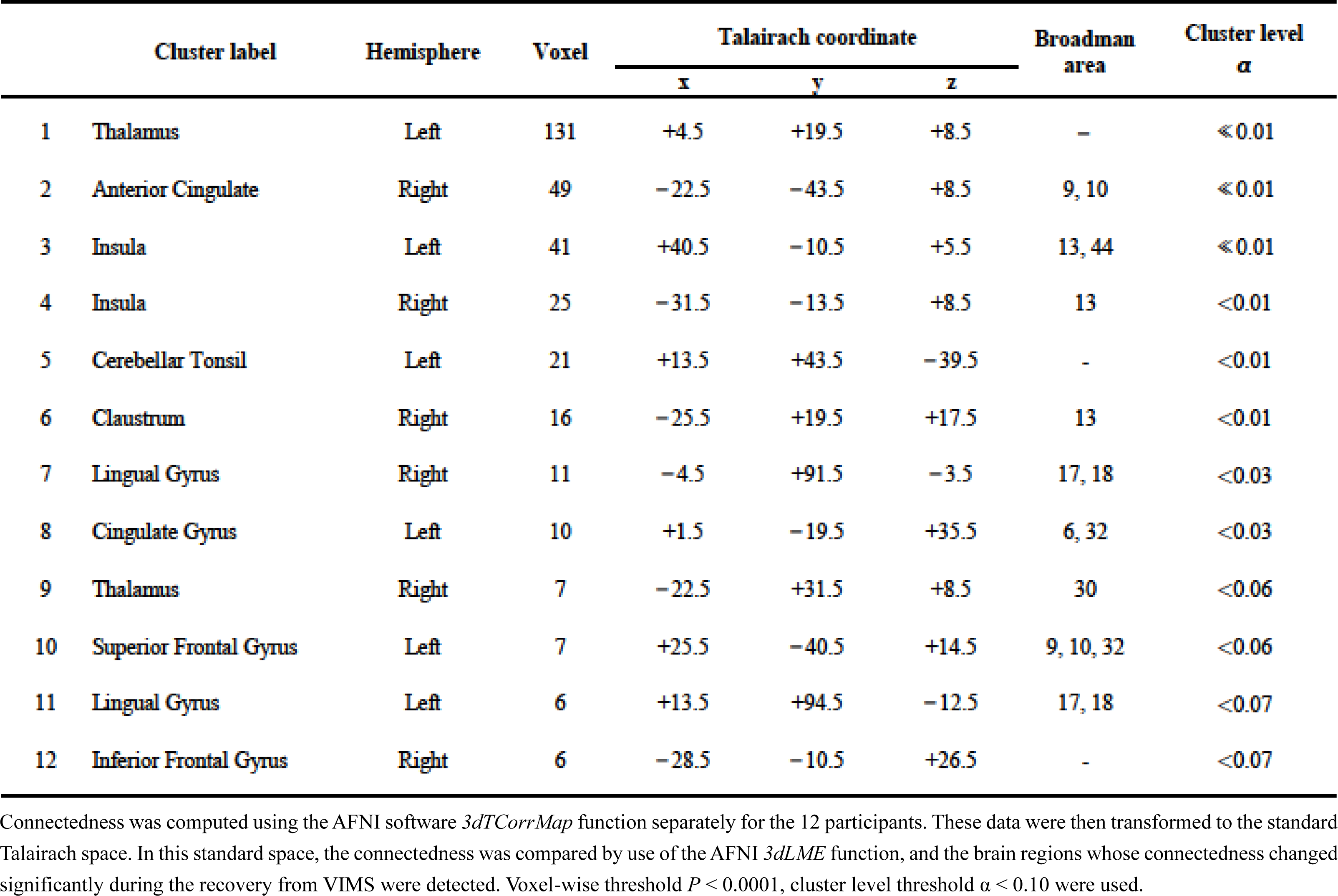
Brain regions whose connectedness increased significantly during the recovery phase from visually induced motion sickness.

**Table 3.** Brain region pairs whose functional connectivity increased during the recovery phase of visually induced motion sickness (VIMS) and the GROUP × PHASE interaction effect.

| Seed label | Cluster label | Hemisphere | GROUP × PHASE interaction |  | Voxel | Talairach coordinate |  |  | Broadman area |
| --- | --- | --- | --- | --- | --- | --- | --- | --- | --- |
| | | | Test statistics<br>*P* value ($\chi^2$) | Effect size<br>$f^2$ | | *x* | *y* | *z* | |
| L Thalamus | 1 Insula | Right | .003 (17.45) | 0.11 | 21 | −34.5 | −13.5 | +2.5 | 13, 47 |
|  | 2 Inferior Parietal Lobule | Right | .014 (14.48) | 0.12 | 9 | −55.5 | +28.5 | +26.5 | 2, 13, 40 |
|  | 3 Declive | Right | .0003 (20.12) | 0.10 | 8 | −4.5 | +58.5 | −12.5 | – |
| L Insula | 4 Thalamus | Left | .001 (18.99) | 0.11 | 15 | +4.5 | +10.5 | +11.5 | – |
|  | 5 Inferior Parietal Lobule | Right | .004 (16.93) | 0.28 | 9 | −58.5 | +28.5 | +29.5 | 2, 40 |
| R Insula | 6 Thalamus | Left | $5 \times 10^{-6}$ (25.78) | 0.15 | 23 | +7.5 | +16.5 | +5.5 | – |
|  | 7 Superior Temporal Gyrus | Left | .002 (18.81) | 0.16 | 13 | +55.5 | +31.5 | +11.5 | 22, 41, 42 |
|  | 8 Inferior Temporal Gyrus | Left | .001 (19.26) | 0.21 | 9 | +49.5 | +58.5 | −9.5 | 19, 20, 37 |
| L Cerebellar Tonsil | 9 Parahippocampal Gyrus | Left | .005 (16.37) | 0.18 | 11 | +16.5 | +16.5 | −15.5 | – |
| R Claustrum | 10 Cingulate Gyrus | Left | .0004 (21.52) | 0.27 | 83 | +1.5 | −22.5 | +26.5 | 24, 32 |
|  | 11 Inferior Parietal Lobule | Right | .002 (18.72) | 0.23 | 22 | −55.5 | +31.5 | +26.5 | 13, 40, 42 |
|  | 12 Inferior Frontal Gyrus | Left | .003 (17.45) | 0.24 | 17 | +40.5 | −25.5 | +2.5 | 13, 45, 47 |
|  | 13 Superior Temporal Gyrus | Left | .0002 (23.07) | 0.21 | 11 | +58.5 | +25.5 | +11.5 | 22, 40, 41, 42 |
|  | 14 Inferior Parietal Lobule | Left | .004 (16.95) | 0.13 | 8 | +55.5 | +34.5 | +23.5 | 13, 40, 42 |
| L Cingulate Gyrus | 15 Insula | Right | $5 \times 10^{-5}$ (25.62) | 0.23 | 19 | −25.5 | +25.5 | +20.5 | 13 |
| L Superior Frontal Gyrus | 16 Medial Frontal Gyrus | Left | $5 \times 10^{-5}$ (25.53) | 0.20 | 67 | +4.5 | −25.5 | +41.5 | 8, 32 |
|  | 17 Middle Temporal Gyrus | Left | .004 (17.16) | 0.30 | 12 | +46.5 | +52.5 | −0.5 | 19, 37 |
|  | 18 Lentiform Nucleus | Left | .0001 (17.72) | 0.29 | 8 | +28.5 | +19.5 | +11.5 | 13 |
| R Inferior Frontal Gyrus | 19 Inferior Frontal Gyrus | Left | .002 (18.80) | 0.31 | 14 | +40.5 | −28.5 | −0.5 | 13, 45, 47 |

**Table 4.** Statistical analysis of the functional connectivity of brain region pairs for the visually induced motion sickness (VIMS) group.

| Seed label | Cluster label | Hemisphere | Rest-1 vs. Rest-2 |  |  | Rest-2 vs. Rest-3 |  |  | Rest-1 vs. Rest-3 |  |  |
| --- | --- | --- | --- | --- | --- | --- | --- | --- | --- | --- | --- |
| | | | Coefficient<br>Estimate ± SE | Test statistics<br>*P* value ($\chi^2$) | Effect size<br>Cohen's *d* | Coefficient<br>Estimate ± SE | Test statistics<br>*P* value ($\chi^2$) | Effect size<br>Cohen's *d* | Coefficient<br>Estimate ± SE | Test statistics<br>*P* value ($\chi^2$) | Effect size<br>Cohen's *d* |
| L Thalamus | 1 Insula | Right | -0.07 ± 0.07 | > 1 (1.05) | 0.42 | 0.34 ± 0.07 | 5×10<sup>-5</sup> (25.70) | 2.07 | 0.27 ± 0.07 | .006 (16.37) | 1.65 |
| | 2 Inferior Parietal Lobule | Right | -0.04 ± 0.06 | > 1 (0.44) | 0.27 | 0.30 ± 0.06 | 4×10<sup>-5</sup> (25.94) | 2.08 | 0.26 ± 0.06 | .001 (19.61) | 1.81 |
| | 3 Declive | Right | -0.03 ± 0.04 | > 1 (0.82) | 0.37 | 0.27 ± 0.04 | 3×10<sup>-12</sup> (57.83) | 3.10 | 0.24 ± 0.04 | 2×10<sup>-9</sup> (44.86) | 2.73 |
| L Insula | 4 Thalamus | Left | 0.02 ± 0.05 | > 1 (0.10) | 0.13 | 0.27 ± 0.05 | 2×10<sup>-6</sup> (32.17) | 2.32 | 0.29 ± 0.05 | 2×10<sup>-7</sup> (35.93) | 2.45 |
| | 5 Inferior Parietal Lobule | Right | -0.04 ± 0.07 | > 1 (0.35) | 0.24 | 0.43 ± 0.07 | 8×10<sup>-8</sup> (38.05) | 2.52 | 0.39 ± 0.07 | 3×10<sup>-6</sup> (31.12) | 2.28 |
| R Insula | 6 Thalamus | Left | -0.06 ± 0.05 | > 1 (1.39) | 0.48 | 0.40 ± 0.05 | 2×10<sup>-11</sup> (53.99) | 3.00 | 0.34 ± 0.05 | 8×10<sup>-8</sup> (38.05) | 2.52 |
| | 7 Superior Temporal Gyrus | Left | 0.11 ± 0.08 | > 1 (2.13) | 0.60 | 0.30 ± 0.08 | .010 (15.40) | 1.60 | 0.41 ± 0.08 | .0003 (29.00) | 2.20 |
| | 8 Inferior Temporal Gyrus | Left | 0.11 ± 0.07 | > 1 (2.79) | 0.68 | 0.26 ± 0.07 | .016 (14.53) | 1.56 | 0.38 ± 0.07 | 5×10<sup>-6</sup> (30.05) | 2.24 |
| L Cerebellar Tonsil | 9 Parahippocampal Gyrus | Left | -0.003 ± 0.08 | > 1 (0.001) | 0.01 | 0.32 ± 0.08 | .008 (15.92) | 1.63 | 0.32 ± 0.08 | .009 (15.65) | 1.62 |
| R Claustrum | 10 Cingulate Gyrus | Left | -0.01 ± 0.06 | > 1 (0.05) | 0.09 | 0.33 ± 0.06 | 3×10<sup>-6</sup> (31.22) | 2.28 | 0.32 ± 0.06 | 9×10<sup>-6</sup> (28.86) | 2.19 |
| | 11 Inferior Parietal Lobule | Right | -0.06 ± 0.07 | > 1 (0.64) | 0.33 | 0.33 ± 0.07 | .0002 (22.58) | 1.94 | 0.27 ± 0.07 | .009 (15.60) | 1.61 |
| | 12 Inferior Frontal Gyrus | Left | -0.004 ± 0.05 | > 1 (0.01) | 0.03 | 0.29 ± 0.05 | 5×10<sup>-6</sup> (30.03) | 2.24 | 0.28 ± 0.05 | 8×10<sup>-6</sup> (29.17) | 2.21 |
| | 13 Superior Temporal Gyrus | Left | 0.11 ± 0.07 | > 1 (2.30) | 0.62 | 0.27 ± 0.07 | .019 (14.19) | 1.54 | 0.38 ± 0.07 | 1×10<sup>-5</sup> (27.91) | 2.16 |
| | 14 Inferior Parietal Lobule | Left | 0.01 ± 0.06 | > 1 (0.01) | 0.04 | 0.26 ± 0.06 | .002 (18.80) | 1.77 | 0.27 ± 0.06 | .001 (19.67) | 1.81 |
| L Cingulate Gyrus | 15 Insula | Right | -0.002 ± 0.07 | > 1 (0.002) | 0.02 | 0.35 ± 0.07 | 1×10<sup>-5</sup> (28.37) | 2.17 | 0.34 ± 0.07 | 1×10<sup>-5</sup> (27.97) | 2.16 |
| L Superior Frontal Gyrus | 16 Medial Frontal Gyrus | Left | 0.04 ± 0.06 | > 1 (0.51) | 0.29 | 0.26 ± 0.06 | .0003 (22.06) | 1.92 | 0.31 ± 0.06 | 7×10<sup>-6</sup> (29.29) | 2.21 |
| | 17 Middle Temporal Gyrus | Left | 0.06 ± 0.07 | > 1 (0.68) | 0.34 | 0.25 ± 0.07 | .090 (11.26) | 1.37 | 0.31 ± 0.07 | .003 (17.45) | 1.71 |
| | 18 Lentiform Nucleus | Left | 0.10 ± 0.07 | > 1 (1.71) | 0.53 | 0.27 ± 0.07 | .026 (13.60) | 1.51 | 0.37 ± 0.07 | 7×10<sup>-5</sup> (24.96) | 2.04 |
| R Inferior Frontal Gyrus | 19 Inferior Frontal Gyrus | Left | 0.001 ± 0.05 | > 1 (0.0004) | 0.01 | 0.24 ± 0.05 | 2×10<sup>-5</sup> (27.66) | 2.15 | 0.24 ± 0.05 | 1×10<sup>-5</sup> (27.87) | 2.16 |

**Table 5.** Statistical analysis of the functional connectivity of brain region pairs for the healthy group.

| Seed label | Cluster label | Hemisphere | Rest-1 vs. Rest-2 |  |  | Rest-2 vs. Rest-3 |  |  | Rest-1 vs. Rest-3 |  |  |
| --- | --- | --- | --- | --- | --- | --- | --- | --- | --- | --- | --- |
| | | | Coefficient<br>Estimate ± SE | Test statistics<br>*P* value ($\chi^2$) | Effect size<br>Cohen's *d* | Coefficient<br>Estimate ± SE | Test statistics<br>*P* value ($\chi^2$) | Effect size<br>Cohen's *d* | Coefficient<br>Estimate ± SE | Test statistics<br>*P* value ($\chi^2$) | Effect size<br>Cohen's *d* |
| L Thalamus | 1 Insula | Right | -0.01 ± 0.05 | > 1 (0.07) | 0.11 | -0.06 ± 0.05 | > 1 (1.25) | 0.46 | -0.07 ± 0.05 | > 1 (1.92) | 0.57 |
| | 2 Inferior Parietal Lobule | Right | -0.06 ± 0.07 | > 1 (0.66) | 0.33 | -0.05 ± 0.07 | > 1 (0.49) | 0.29 | -0.11 ± 0.07 | > 1 (2.30) | 0.62 |
| | 3 Declive | Right | -0.04 ± 0.06 | > 1 (0.59) | 0.31 | -0.05 ± 0.06 | > 1 (0.82) | 0.37 | -0.10 ± 0.06 | > 1 (2.79) | 0.68 |
| L Insula | 4 Thalamus | Left | -0.07 ± 0.07 | > 1 (1.13) | 0.43 | -0.06 ± 0.07 | > 1 (0.90) | 0.39 | -0.14 ± 0.07 | > 1 (4.05) | 0.82 |
| | 5 Inferior Parietal Lobule | Right | -0.11 ± 0.07 | > 1 (2.30) | 0.62 | -0.04 ± 0.07 | > 1 (0.24) | 0.20 | -0.07 ± 0.07 | > 1 (1.05) | 0.42 |
| R Insula | 6 Thalamus | Left | -0.02 ± 0.05 | > 1 (0.13) | 0.15 | -0.04 ± 0.05 | > 1 (0.88) | 0.38 | -0.06 ± 0.05 | > 1 (1.70) | 0.53 |
| | 7 Superior Temporal Gyrus | Left | 0.02 ± 0.06 | > 1 (0.08) | 0.11 | -0.09 ± 0.06 | > 1 (2.25) | 0.61 | -0.07 ± 0.06 | > 1 (1.50) | 0.50 |
| | 8 Inferior Temporal Gyrus | Left | -0.05 ± 0.05 | > 1 (1.03) | 0.41 | -0.04 ± 0.05 | > 1 (0.45) | 0.27 | -0.09 ± 0.05 | > 1 (2.84) | 0.69 |
| L Cerebellar Tonsil | 9 Parahippocampal Gyrus | Left | -0.05 ± 0.07 | > 1 (0.45) | 0.27 | -0.10 ± 0.07 | > 1 (2.12) | 0.59 | -0.15 ± 0.07 | > 1 (4.51) | 0.87 |
| R Claustrum | 10 Cingulate Gyrus | Left | -0.03 ± 0.05 | > 1 (0.41) | 0.26 | -0.06 ± 0.05 | > 1 (1.29) | 0.46 | -0.09 ± 0.05 | > 1 (3.15) | 0.72 |
| | 11 Inferior Parietal Lobule | Right | 0.01 ± 0.06 | > 1 (0.06) | 0.10 | -0.12 ± 0.06 | > 1 (4.20) | 0.84 | -0.11 ± 0.06 | > 1 (3.26) | 0.74 |
| | 12 Inferior Frontal Gyrus | Left | -0.08 ± 0.06 | > 1 (1.63) | 0.52 | -0.03 ± 0.06 | > 1 (0.25) | 0.20 | -0.11 ± 0.06 | > 1 (3.14) | 0.72 |
| | 13 Superior Temporal Gyrus | Left | -0.08 ± 0.04 | > 1 (4.84) | 0.90 | -0.04 ± 0.04 | > 1 (1.06) | 0.42 | -0.11 ± 0.04 | .142 (10.43) | 1.32 |
| | 14 Inferior Parietal Lobule | Left | -0.07 ± 0.05 | > 1 (2.00) | 0.58 | -0.04 ± 0.05 | > 1 (0.46) | 0.27 | -0.11 ± 0.05 | > 1 (4.34) | 0.85 |
| L Cingulate Gyrus | 15 Insula | Right | -0.06 ± 0.04 | > 1 (2.55) | 0.65 | -0.06 ± 0.04 | > 1 (2.69) | 0.67 | -0.12 ± 0.04 | .137 (10.49) | 1.32 |
| L Superior Frontal Gyrus | 16 Medial Frontal Gyrus | Left | 0.003 ± 0.03 | > 1 (0.02) | 0.06 | -0.08 ± 0.03 | .313 (8.97) | 1.22 | -0.08 ± 0.03 | .502 (8.11) | 1.16 |
| | 17 Middle Temporal Gyrus | Left | -0.06 ± 0.05 | > 1 (1.33) | 0.47 | -0.06 ± 0.05 | > 1 (0.70) | 0.53 | -0.12 ± 0.05 | > 1 (6.04) | 1.00 |
| | 18 Lentiform Nucleus | Left | -0.04 ± 0.06 | > 1 (0.50) | 0.29 | -0.07 ± 0.06 | > 1 (1.15) | 0.44 | -0.11 ± 0.06 | > 1 (3.18) | 0.73 |
| R Inferior Frontal Gyrus | 19 Inferior Frontal Gyrus | Left | -0.02 ± 0.05 | > 1 (0.14) | 0.15 | -0.07 ± 0.05 | > 1 (1.70) | 0.53 | -0.09 ± 0.05 | > 1 (2.81) | 0.68 |

The description is the same as that of Table 4 but for the healthy participant group.

Among the three SSQ subscales, the Disorientation score was similar to the Total Score (PHASE × GROUP interaction: χ^2^(2) = 19.16, *P* = 0.0002; *post hoc* test: the VIMS group: Rest-1 *vs*. Rest-2: χ^2^(2) = 0.02, *P* > 1; Rest-2 *vs*. Rest-3: χ^2^(2) = 18.52, *P* = 0.0002; Rest-1 *vs*. Rest-3: χ^2^(2) = 19.87, *P* = 0.0001; the healthy group: Rest-1 *vs*. Rest-2: χ^2^(2) = 0.53, *P* > 1; Rest-2 *vs*. Rest-3: χ^2^(2) < 0.01, *P* > 1; Rest-1 *vs*. Rest-3: χ^2^(2) = 0.52, *P* > 1, Bonferroni corrected). As for Nausea and Oculomotor, the PHASE × GROUP interactions were not significant (Nausea, χ^2^(2) = 7.74, *P* = 0.084; Oculomotor, χ^2^(2) = 5.48, *P* = 0.259; Bonferroni corrected), and the main effect of PHASE was also not significant (Nausea, χ^2^(2) = 4.28, *P* = 0.539; Oculomotor, χ^2^(2) = 4.62, *P* = 0.444; Bonferroni corrected).

### Brain connectedness mapping

To test our hypothesis that the recovery phase of VIMS induces changes in functional connectivity in some brain regions, we examined the substantial changes in functional connectivity by calculating connectedness, which is the mean correlation coefficient of a specific voxel time-series to other voxel time-series (Gotts *et al*., 2012), and then by mapping the results in Talairach space. As anticipated, the map revealed 12 brain regions with significant increases in connectedness during Rest-3 in the VIMS group (Table 2 and Fig. 3). The 12 regions included the primary visual cortex in the occipital cortex (the left and right lingual gyrus; Fig. 3k and g, respectively) and the left and right insula (Fig. 3c and d) and cingulate regions (the right anterior cingulate and left cingulate gyrus; Fig. 3b and h, respectively). In addition, the left superior frontal gyrus (Fig. 3j) and the right inferior frontal gyrus (Fig. 3l) were included in the 12 brain regions. Furthermore, connectedness significantly increased in the cerebellum (left cerebellar tonsil; Fig. 3e) and the subcortical region (left and right thalamus and the right claustrum; Fig. 3a, i, and f, respectively) during the recovery phase of VIMS. In contrast, no brain region had a significant decrease in connectedness during the recovery phase.

Each row is as follows: a, left thalamus; b, anterior cingulate; c, left insula; d, right insula; e, cerebellar tonsil; f, claustrum; g, right lingual gyrus; h, cingulate gyrus; i, right thalamus; j, left superior frontal gyrus; k, left lingual gyrus; and l, inferior frontal gyrus. Each column shows, from left to right, axial, sagittal, and coronal brain images. The following thresholds were set: voxel-wise threshold *P* < 0.0001, cluster level threshold *α* < 0.10.

### ROI-based functional connectivity analyses

For the 12 ROIs with increased connectedness, the question arose as to whether functional connectivity had increased in a broad area or in only specific regions. To address this question, the entire brain was screened for functional connectivity using the 12 ROIs as seeds. It was found that 8 of the 12 seed regions showed increases in functional connectivity with specific regions during only Rest-3 (recovery phase) for the VIMS group. These regions were the left thalamus, left and right insula, left cerebellar tonsil, right claustrum, left cingulate gyrus, left superior frontal gyrus, and left inferior frontal gyrus. Each of these regions had such selective increases in functional connectivity with several other regions, resulting in the following 19 region pairs: left thalamus - right insula, left thalamus - right inferior parietal lobule, left thalamus - right declive, left insula - left thalamus, left insula - left inferior parietal lobule, right insula - left superior temporal gyrus, right insula - left inferior temporal gyrus, left cerebellar tonsil - left parahippocampal gyrus, right claustrum - left cingulate gyrus, right claustrum - right inferior parietal lobule, right claustrum - left inferior frontal gyrus, right claustrum - left superior temporal gyrus, right claustrum - left inferior parietal lobule, left cingulate gyrus - right insula, left superior frontal gyrus - left medial frontal gyrus, left superior frontal gyrus - left middle temporal gyrus, left superior frontal gyrus - left lentiform nucleus, and right inferior frontal gyrus - left inferior frontal gyrus (Table 3). The locations of the 19 pairs are shown in the left panels of Figure 4 and Supplementary Figure 1. The other 4 of the 12 seed regions had no such selective changes in functional connectivity.

The phase (Rest-1, -2, and -3) × group (Healthy and VIMS) interaction effect can be clearly seen in the right panels of Figure 4 and Supplementary Figure 1. The functional connectivity of the VIMS group (yellow solid line) remained comparable in Rest-1 and Rest-2 but increased in Rest-3 (recovery phase), whereas that of the healthy group (white broken line) did not show an increase in any phase. These results were statistically confirmed by a linear mixed-effects model analysis. The interaction effects of resting phases (PHASE: Rest-1, Rest-2, and Rest-3) and participants’ group (GROUP: VIMS and healthy) were statistically significant for all 19 pairs (Table 3). We thus analysed the simple effect of PHASE separately for the VIMS and healthy groups. The statistical results for the VIMS and healthy groups are shown in Tables 4 and 5, respectively. The selective increase in functional connectivity for the VIMS group during Rest-3 (recovery phase) were statistically significant for almost all pairs and the effect sizes (Cohen’s *d*) were relatively large (Table 4). For example, for the right insula - left thalamus pair (#6 in Tables 4 and 5), the simple effect of the Rest-3 PHASE was statistically significant for only the VIMS group (Rest-2 *vs*. Rest-3: χ^2^(2) = 53.99, *P* = 2 × 10^−11^, Cohen’s *d* = 3.00; Rest-1 *vs*. Rest-3: χ^2^(2) = 38.05, *P* = 8 × 10^−8^, Cohen’s *d* = 2.52, Bonferroni corrected) (#6 in Table 4), whereas this effect was not found for the other control conditions (VIMS group: Rest-1 *vs*. Rest-2: χ^2^(2) = 1.39, *P* > 1, Cohen’s *d* = 0.48; healthy group: Rest-1 *vs*. Rest-2: χ^2^(2) = 0.13, *P* > 1, Cohen’s *d* = 0.15; Rest-2 *vs*. Rest-3: χ^2^(2) = 0.88, *P* > 1, Cohen’s *d* = 0.38; Rest-1 *vs*. Rest-3: χ^2^(2) = 1.70, *P* > 1, Cohen’s *d* = 0.53, Bonferroni corrected) (#6 in Tables 4 and 5). Similar selective changes in functional connectivity were observed in the other 18 region pairs (see Tables 3, 4, and 5). However, as an exception, the functional connectivity between the left superior frontal gyrus and left middle temporal gyrus showed a tendency toward an increase in the second *vs*. third rest phases in the VIMS group (#17 in Table 4).

### Correlation analyses

To determine whether the increases in functional connectivity were associated with subjective recovery from VIMS, we conducted correlation analyses between the change in functional connectivity and that in the SSQ Total Scores for the 19 pairs of brain regions (Fig. 5). The following 5 pairs showed statistically significant negative correlations, indicating regions in which greater improvements were associated with greater recovery from VIMS: the right insula–left superior temporal gyrus (Spearman’s rank correlation ρ = −0.83, *P* = 0.017, Fig. 5a), right claustrum–right inferior parietal lobule (ρ = – 0.83, *P* = 0.016, Fig. 5b), right claustrum–left superior temporal gyrus (ρ = –0.92, *P* = 0.0004, Fig. 5c), right claustrum–left inferior parietal lobule (ρ = –0.80, *P* = 0.038, Fig. 5d), and left superior frontal gyrus–left lentiform nucleus (ρ = –0.80, *P* = 0.038, Fig. 5e, all Bonferroni corrected). The correlation was not statistically significant in the remaining 14 pairs: the left thalamus–right insula (ρ = −0.64, *P* = 0.469), left thalamus–right inferior parietal lobule (ρ = −0.63, *P* = 0.510), left thalamus–right declive (ρ = −0.71, *P* = 0.181), left insula–left thalamus (ρ = −0.73, *P* = 0.137), left insula–left inferior parietal lobule (ρ = −0.62, *P* = 0.576), right insula–left thalamus (ρ = −0.67, *P* = 0.330), right insula–left inferior temporal gyrus (ρ = −0.61, *P* = 0.647), left cerebellar tonsil–left parahippocampal gyrus (ρ = −0.71, *P* = 0.181), right claustrum–left cingulate gyrus (ρ = –0.75, *P* = 0.095), right claustrum–left inferior frontal gyrus (ρ = −0.74, *P* = 0.115), left cingulate gyrus–right insula (ρ = −0.70, *P* = 0.212), left superior frontal gyrus– left medial frontal gyrus (ρ = −0.63, *P* = 0.510), left superior frontal gyrus–left middle temporal gyrus (ρ = −0.58, *P* = 0.932), and right inferior frontal gyrus–left inferior frontal gyrus (ρ = −0.61, *P* = 0.698, all Bonferroni corrected).

**Figure 5.**
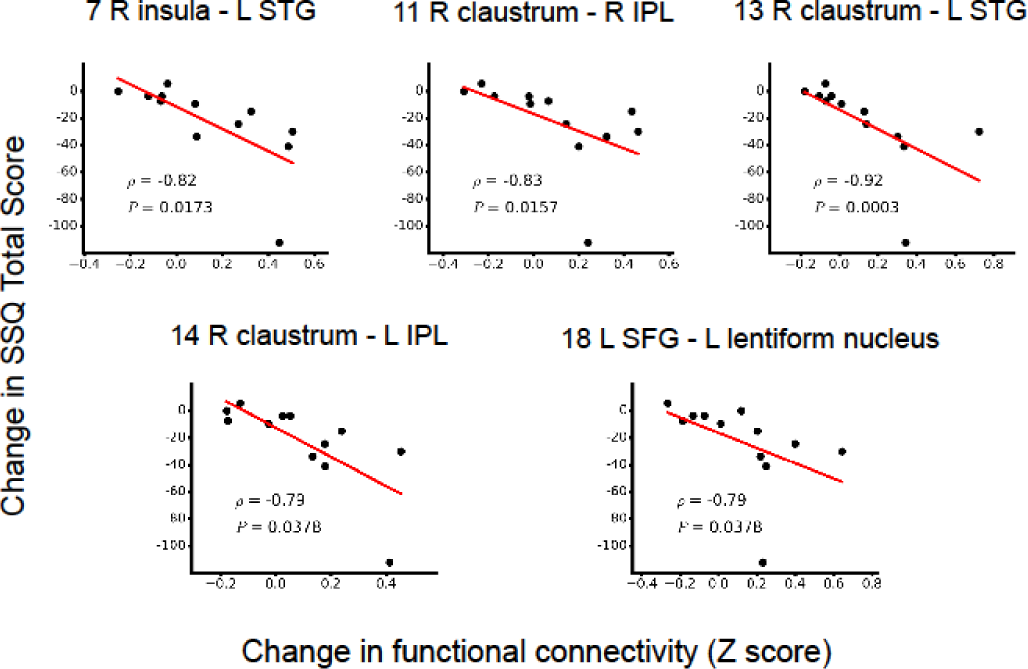
Correlation plots between the change in the Total Score (TS) of the Simulator Sickness Questionnaire (SSQ) and the change in functional connectivity. The vertical axis is the change in the TS of the SSQ: Rest-3 minus the average of Rest-1 and Rest-2; the more VIMS symptoms recovered, the more this score decreases. The horizontal axis is the change in functional connectivity: Rest-3 minus the average of Rest-1 and Rest-2. In the graphs, ρ indicates Spearman’s rank correlation coefficient, with the corresponding *P* value shown below. The *P* values were Bonferroni corrected for multiple comparisons among the 19 brain region pairs. Abbreviations: IPL, inferior parietal lobule; L, left; R, right; SFG, superior frontal gyrus; STG, superior temporal gyrus.

### Global network analyses

The above connectedness map shows the 12 brain regions with recovery-selective increases in connectedness (Fig.3 and Table 2). To explore the possibility that these regions constituted larger functional networks, we conducted a multivariate analysis in which the network of these regions was analysed based on distance matrices representing dissimilarity among the fMRI time-series in each pair of these regions (Supplementary Figure 2), from which hierarchical clustering dendrograms were derived. Figure 6 compares the results of the VIMS group (a, c) with those of the healthy group (b, d). As clearly shown in Fig. 6, the 12 brain regions of the VIMS group showed more global network structures for both recovery (Rest-3) and non-recovery (the average of Rest-1 and Rest-2) phases, in contrast to the healthy group, for which there was only a statistically significant cluster (marked with asterisks) of the left and right lingual gyri. Notably, the statistically significant clusters of the VIMS group (marked with asterisks) comprised the brain regions with recovery-selective increases in ROI-based functional connectivity (marked with underlines) and the regions with correlation with the SSQ (marked with italics), corroborating the above analyses (Figs. 4 and 5). During the recovery phase (Rest-3), the left thalamus additionally participated in the visual cortical cluster, and this cluster was combined with the other cluster of the left/right insular and cingulate areas in a higher level. Interestingly, during the phases before sickness developed, the VIMS group already showed a distinct cluster composed of the left cingulate gyrus and right thalamus in conjunction with the visual cortical cluster. Some of these networks were also independently confirmed by dictionary learning-based connectivity mapping (Supplementary Figure 3).

**Figure 6.**
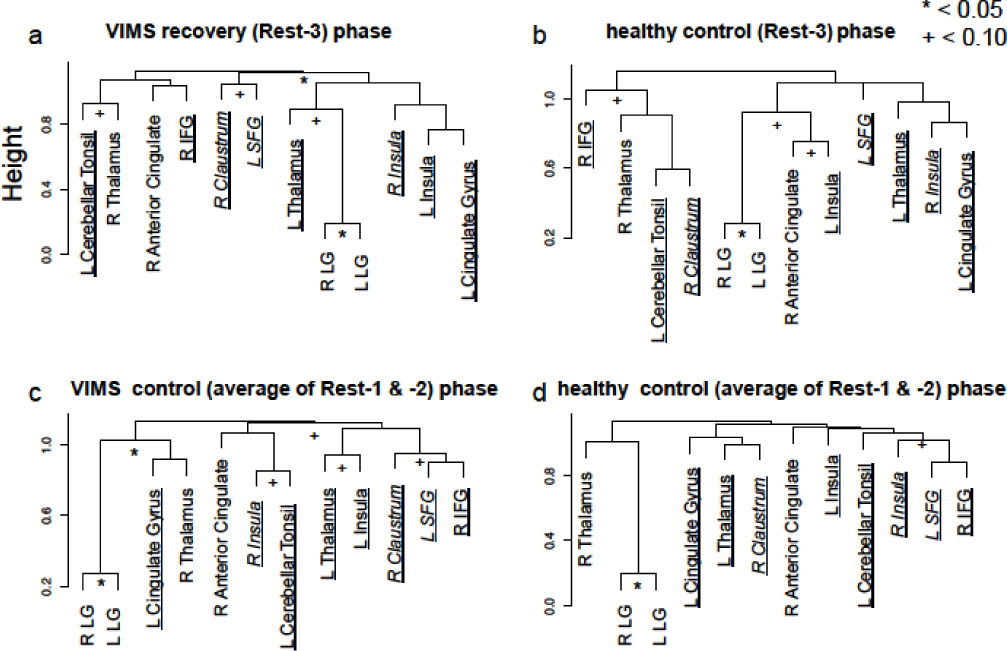
Hierarchical clustering trees of the 12 brain regions showing recovery-selective increases in connectedness. (a) The dendrogram of the VIMS group for the recovery phase (Rest-3), which was derived from the distance matrix (Supplementary Fig. 2a), whose elements indicate 1 minus absolute partial correlation between each pair of the 12 regions. Statistically significant clusters are marked with an asterisk (*P* < 0.05) or plus (*P* < 0.1). Brain regions showing recovery-selective increases in the ROI-based functional connectivity and correlations with SSQ are underlined and in italics, respectively. (b) The same as (a) but for the healthy group. (c) The same as (a) but for the control phase (the average of Rest-1 and Rest-2). (d) The same as (c) but for the healthy group. Abbreviations: L, left; R, right; LG, lingual gyrus; SFG, superior frontal gyrus; IFG, inferior frontal gyrus.

## Discussion

In this study, we investigated resting-state fMRI functional connectivity during the recovery phase of VIMS. The analysis showed that the functional connectivity was increased in some brain areas, including the insula, cingulate, and visual cortical regions. These regions were also found to constitute large interactive networks. Furthermore, some of the increases in connectivity were strongly correlated with the subjective awareness of recovery from VIMS (decrease in subjective SSQ scores) for the following 5 pairs of brain regions: insula–superior temporal gyrus, claustrum–left inferior parietal lobule, claustrum–right inferior parietal lobule, claustrum–superior temporal gyrus, and superior frontal gyrus–lentiform nucleus. Although the sample size was not large (12 participants), to the best of our knowledge, this is the first study to elucidate the brain process in the recovery phase of VIMS. The above brain regions detected in our study included key regions related to the emergence and evolution of VIMS. The insula had increased connectivity with the cingulate gyrus during recovery from VIMS (Supplementary Figure 1k). The insular and cingulate regions have also shown a correlated activation during a higher nausea state (Napadow *et al*., 2013). Additionally, the insula showed a significant correlation between sympathovagal balance and the connectivity with a middle temporal region (MT+/V5) during a visually induced nausea sensation (Toschi *et al*., 2017), which is a visual motion-sensitive area. In addition, the middle temporal region was reported to decrease the inter-hemispheric synchronization during VIMS (Miyazaki *et al*., 2015). This key area had increased connectivity with the superior frontal region during recovery from VIMS (Supplementary Figure 1m). In addition to these key brain areas, several regions detected in our analysis of the recovery from VIMS, such as the inferior frontal gyrus (Supplementary Figure 1j and 1N), cerebellar tonsil (Supplementary Figure 1h), and declive (Supplementary Figure 1c), have been suggested to be associated with the evolution of VIMS (Farmer *et al*., 2015). These results thus conformed with our initial expectations that key brain regions related to the emergence and evolution of VIMS also play roles in the recovery from VIMS.

Closer looks at the changes in functional connectivity along the resting phases Rest-1, Rest-2, and Rest-3 (Fig. 4) suggest two different groups among the 5 pairs of brain regions that may have distinct roles. For the first group, the right insula–superior temporal gyrus (Fig. 4a) and the superior frontal gyrus–lentiform nucleus (Fig. 4e), functional connectivity remained roughly comparable in the healthy and VIMS groups before the experience of VIMS (Rest-1 and Rest-2) but increased only in the VIMS group during recovery from VIMS (Rest-3), indicating temporary reorganisations of neural circuitry triggered by the experience of unusual motion environments. In contrast, for the second group, the claustrum–inferior parietal lobule (Fig. 4b and 4d), functional connectivity was weaker in the VIMS group than in the healthy group before the onset of VIMS but became comparable in the two groups during recovery from VIMS. A similar intrinsic difference between the groups was also observed in the dendrogram (Fig. 6c vs. Fig. 6d). These results imply that VIMS group participants might have weaker connectivities between specific brain regions that reflect VMIS susceptibility and/or tolerance. It would be interesting to address MS susceptibility and/or tolerance in a future study with a larger sample size.

Why did the functional connectivity between the brain regions increase during recovery from VIMS? Although it is impossible to explain the precise mechanisms underlying the results, it is tempting to discuss the mechanisms by considering our results in conjunction with the collective knowledge in the field. Here we propose three possibilities and discuss them in relation to current understanding of VIMS and the related hypotheses. The first possibility is that the functional connectivity changes observed during the recovery phase would be associated with interoceptive awareness (perception of one’s own bodily state) because interoceptive awareness is expected to increase to alleviate the symptoms of VIMS, such as dizziness, eye pain, and nausea. In this regard, it is interesting that the connectivities, including insular and cingulate regions, were increased in our study. The insular cortex, the centre of the neural basis of interoception, is also regarded as the limbic sensory cortex (Craig, 2002; Craig & Craig, 2009) because this brain site is involved in the homeostatic regulation of the brain stem. In addition, the cingulate gyrus is known as the limbic motor cortex, which plays an important role in interoception together with the insular cortex (Craig, 2002; Craig & Craig, 2009). Indeed, according to Napadow *et al*. (2013), in association with the insular cortex, the cingulate cortex is coactivated when the perception of nausea is triggered by visual stimuli. It has been pointed out that the claustrum, which had functional connectivity with multiple brain regions in this study, has been speculated to be involved in the processing of conscious perception, suggesting its association with interoceptive perception (Crick & Koch, 2005). Therefore, the strengthened brain networks centring on the insula and claustrum in this study may reflect the internal perception of a poor physical state triggered by VIMS. Similarly, after observing the strengthening of the amygdala network (including the insular cortex) immediately after the induction of psychological stress by visual stimuli, van Marle *et al*. (2009) suggested that the strengthened amygdala network reflects negative hypervigilance.

The second possibility is that the changes in functional connectivity observed during the recovery phase of VIMS would reflect plastic changes in the neural circuit related to visual processing such as visual attention and eye movement control. In this regard, we want to note that visual processing regions, including the middle temporal gyrus, which is a visual motion-sensitive area, had a significant increase in functional connectivity during the recovery phase of VIMS in our study. Indeed, the middle temporal gyrus had a strong connectivity with the superior frontal gyrus (Supplementary Figure 1m and #17 in Tables 3, 4, and 5). Interestingly, similar results were reported in previous neuroimaging studies on visual or motor learning (Albert *et al*., 2009a, b; Lewis *et al*., 2009; Stevens *et al*., 2009, 2015). Lewis *et al*. (2009) measured resting-state functional connectivity after a visual shape recognition task and found increased connectivity between the occipital visual area and frontoparietal region, which is involved in visual attention. In addition, Stevens *et al*. (2009) showed that functional connectivities between the neural network in the occipital face and scene areas and frontal cortex were increased during the rest phase immediately after a visual face and scene recognition task and that subsequent memory retrieval performance was predicted by the extent of the changes in the functional connectivity. This suggests that a functional connectivity increase during a rest phase after a task reflects the neuroplastic change needed to improve the efficacy of task processing. In addition to this change in connectivity between occipital visual and frontal regions, brain areas showing increased functional connectivity in this study (the insula–superior temporal gyrus [Figure 4A], insula–inferior temporal gyrus [Supplementary Figure 1g], cingulate gyrus–insula [Supplementary Figure 1k], superior frontal gyrus–medial frontal gyrus [Supplementary Figure 1l], and left and right inferior frontal gyrus [Supplementary Figure 1n]) were coactivated when regulating visual attention or eye movement in previous fMRI studies (Corbetta *et al*., 1998; Dieterich *et al*., 1998, 2003; Konen *et al*., 2005). As for the claustrum having increased connectivity to multiple brain regions in this study, this area plays an important role in visual attention as well as multisensory attention and is activated at the same time as the inferior parietal lobule and interior frontal gyrus (Vohn *et al*., 2007), both of which were detected in this study. Attention is closely associated with eye movement (Corbetta *et al*., 1998), and eye movement has been proposed to cause MS including VIMS in the eye movement hypothesis (Ebenholtz, 1992; Ebenholtz *et al*., 1994). The eye movement hypothesis posits that MS is caused by abnormal eye movement induced in a certain kind of motion environment. Given this prior knowledge and our results, we speculate that the increase in functional connectivity observed during the recovery from VIMS might reflect a plastic change in neural circuits related to visual processing such as visual attention and/or eye movement control.

The supposed neural plasticity might be linked to the sensory conflict hypothesis (Reason & Brand, 1975; Reason, 1978; Oman, 1990), which is a major hypothesis for the occurrence of MSs such as VIMS. This hypothesis suggests that a conflict or mismatch among multiple sensory inputs, including visual and vestibular inputs, would cause MS. Interestingly, this theory also indicates that a conflict not only among sensory inputs, but also between a sensory input and an empirically acquired sensory memory (so-called ‘neural store’) would cause MS. As for this perspective, individuals with MS such as VIMS develop adaptation, and tolerance increases when similar stimuli are repeated because of the reduced conflict between a sensory input and ‘neural store’, which is the memory acquired from the repeated sensory experience (Reason & Brand, 1975; Reason, 1978; Oman, 1990). Therefore, we speculate that an adaptation-related neuroplastic change might occur in the corresponding neural circuits immediately after the induction of VIMS. The neuronal circuit of VIMS and other types of MS is thought to be the functional network centring on visual and vestibular systems. Previous neuroimaging studies of the visual and vestibular responses to stimuli suggested close cooperation between the motion-sensitive visual cortices and vestibular cortex for self-motion perception (Brandt *et al*., 1998; Kleinschmidt *et al*., 2002; Smith *et al*., 2012; Frank *et al*., 2014, 2016). In addition, in the present study, significant changes in functional connectivity were observed in the higher order visual areas of the inferior and middle temporal gyri during recovery from VIMS (Supplementary Figure 1g and 1m; Tables 3, 4, and 5). However, no change in connectivity was observed in the parieto-insular vestibular cortex known as the centre of the neural basis of vestibular sensation. This might have been because visual inputs were the only trigger of VIMS and because the visual system was the sole centre of brain dynamics during the recovery phase.

Finally, the third possibility is that the increased functional connectivity in these brain regions during recovery from VIMS might reflect the maintenance of spatial memory. In our results, the connectivity between the cerebellar tonsil and parahippocampal gyrus increased (Supplementary Figure 1h). The parahippocampal gyrus is said to be the centre of visual spatial information processing (Epstein *et al*., 1998; Epstein, 2008), playing an important role in the recognition and memorisation of the space where one exists. Previous resting-state fMRI studies of memory maintenance have reported that the functional connectivity between the brain regions involved in memory processing significantly increases after memory tasks (Tambini *et al*., 2010; Gordon *et al*., 2014). For example, in Tambini *et al*. (2010), the resting-state connectivity between the hippocampus and lateral occipital complex increased after associative memory tasks, which enabled the prediction of subsequent memory performance. Therefore, it might be possible that the increased connectivity in these regions reflects novel spatial experiences in memory processing in the present participants with VIMS.

If so, our findings might provide some insight into the neural basis of the sensory conflict hypothesis (Reason & Brand, 1975; Reason, 1978; Oman, 1990) mentioned above. The formation of visual spatial memory is extremely important when explaining the adaptation of VIMS and other types of MS. By incorporating the concept of a neural store, the sensory conflict hypothesis explains that MS develops due to a neural mismatch between the accumulated spatial memory (that is, neural store) through experiences and the sensory inputs from the sensory organs. Although the ‘neural store’ has been just a concept, the actual brain network associated with this concept might have been revealed by our findings. If so, these findings can advance our understanding of the neural mechanisms underlying the development and adaptation of MS.

## Acknowledgements

This study was supported by Grants-in-Aid for Scientific Research on Innovative Areas “Shitsukan” (23135517, 25135720) from the Ministry of Education, Culture, Sports, Science and Technology of Japan and a Grant-in-Aid for Scientific Research (16K13506) from the Japan Society for the Promotion of Science (JSPS) to H. Yamamoto.

## Author Contributions

J.M.: Conceptualization, Formal analysis, Investigation, Writing - Original Draft, Writing - Review & Editing.

H.Y.: Project Administration, Supervision, Conceptualization, Methodology, Writing - Original Draft, Writing - Review & Editing, Funding acquisition.

Y.I.: Supervision, Writing - Review.

H.Y., T.Y.: Methodology, Software support, Writing - Review.

T.M., M.U., T.H.: Methodology, Imaging, Writing - Review.

## Data Accessibility

Data in this study are available upon request. Please contact the corresponding author for access.

