## Supplementary Information for "Resting-state functional connectivity predicts recovery from visually induced motion sickness"

**Brain regions whose functional connectivity with the seed regions increased in the recovery phase from visually induced motion sickness (VIMS).**

For 14 brain region pairs presented in Tables 3-5, which was not included in Figure 4, the location in the brain and temporal transition of functional connectivity are shown below.

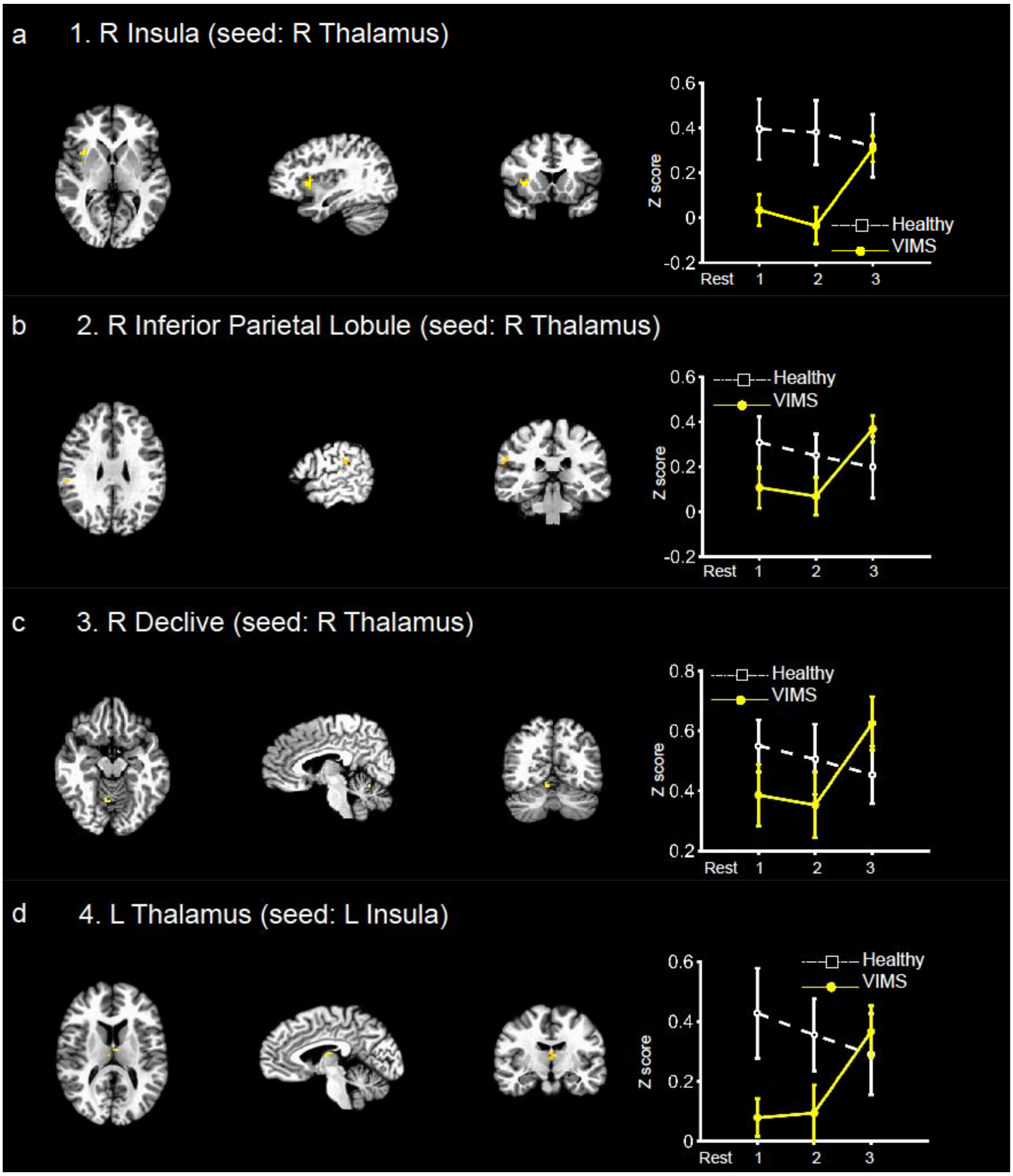

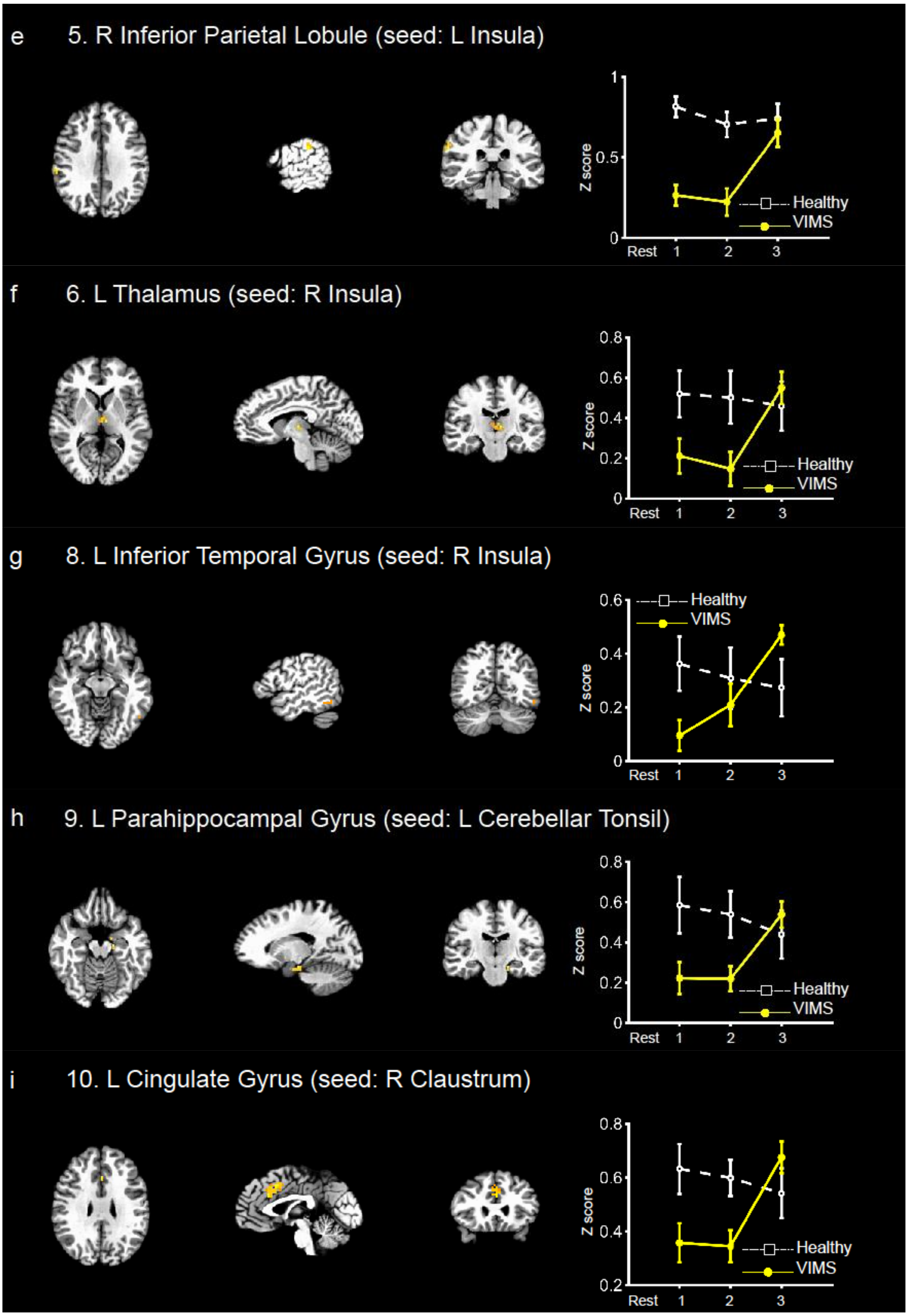

e
5. R Inferior Parietal Lobule (seed: L Insula)
Z score
1
0.5
0
Rest 1 2 3
Healthy
VIMS
f
6. L Thalamus (seed: R Insula)
Z score
0.8
0.6
0.4
0.2
0
Rest 1 2 3
Healthy
VIMS
g
8. L Inferior Temporal Gyrus (seed: R Insula)
Z score
0.6
0.4
0.2
0
Rest 1 2 3
Healthy
VIMS
h
9. L Parahippocampal Gyrus (seed: L Cerebellar Tonsil)
Z score
0.8
0.6
0.4
0.2
0
Rest 1 2 3
Healthy
VIMS
i
10. L Cingulate Gyrus (seed: R Claustrum)
Z score
0.8
0.6
0.4
0.2
Rest 1 2 3
Healthy
VIMS

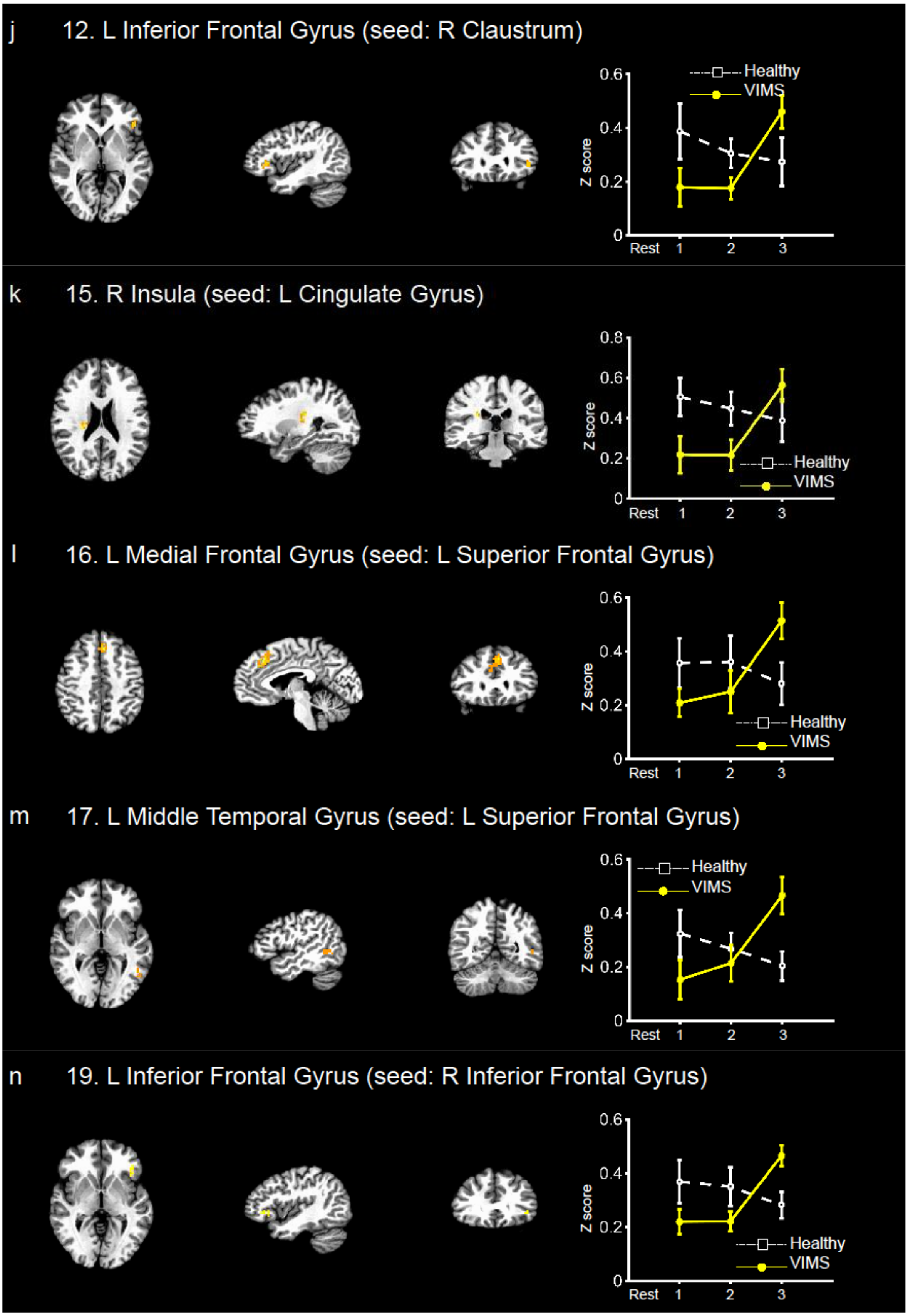


**Supplementary Figure 1. Brain regions whose functional connectivity with the seed**

**regions increased in the recovery phase from visually induced motion sickness (VIMS).**

The effect of head motion (maximum framewise displacement [FD]) on functional connectivity was statistically assessed by using a linear mixed-effects model, in which a within-participants fixed effect of FD and interactions of FD × PHASE and FD × GROUP were included, in addition to the fixed effects of PHASE, GROUP, and PHASE × GROUP and a random effect of each individual participant. If the head movement-related interactions (i.e., FD × PHASE and FD × GROUP) were statistically significant, the statistics for FD were omitted because it was difficult to determine the effect.

### Statistical assessment of the effects of head motion on functional connectivity

To clarify whether head motion affects functional connectivity, we statistically assessed the effect of head movement on functional connectivity. As a measure of head movement, the framewise displacement (FD) (Power et al. 2012) was computed. Then, the maximum value of FD during each rest phase was derived, and the effect of this maximum FD on the functional connectivity of 19 brain region pairs (listed in Tables 3–5) was tested with a linear mixed-effects model analysis. The linear mixed-effects model had fixed effects of the maximum FD (hereafter "FD") in addition to PHASE (Rest-1, -2, and -3) and GROUP (VIMS and healthy group) and three interactions of FD × PHASE, FD × GROUP, and PHASE × GROUP, as well as a random intercept for participants. A standard model selection technique was conducted using the *step* function in the lmerTest package (Kuznetsova *et al.* 2017) of R software. The *step* function statistically tests fixed effects based on the Satterthwaite method and removes ineffective terms from the model.
All 19 brain regions again showed a significant PHASE × GROUP interaction, confirming our results in the main text. The results for the effect of FD are shown in Supplementary Table 1, in which *P* values were not adjusted. For 15 brain region pairs among the 19 pairs, three head motion-related terms (that is the fixed effect of FD and the interactions of FD × PHASE and FD × GROUP) were not statistically significant, suggesting no effect of head motion on functional connectivity. A statistically significant FD × GROUP interaction was obtained for the remaining four pairs: the right claustrum–right inferior parietal lobule, right claustrum–left superior temporal gyrus, right claustrum–left inferior parietal lobule, and left superior frontal gyrus–left medial frontal gyrus. The PHASE × GROUP interaction was again statistically significant, even when the FD × GROUP interaction term and the simple effect of FD were included in the model as covariants. The *post hoc* interaction analyses essentially showed the same results for the VIMS group as those obtained with the original model without the covariants. For the healthy group, a slightly different result was obtained. There were simple effects of PHASE not found by the original model for two of the four pairs: the right claustrum–right inferior parietal lobule and the left superior frontal gyrus–left medial frontal gyrus. For these pairs, recovery-selective decreases in functional connectivity were shown for Rest-3 compared with Rest-2.

**Supplementary Table 1. Statistical analysis of the effect of head movement on functional connectivity.**

| Seed region | | Paired region | Hemispgere | Test statistics<br>*P* value (*F* value) | | |
|---|---|---|---|---|---|---|
| | | | | FD | FD × GROUP | FD × PHASE |
| L Thalamus | 1 | Insula | Right | .739 (.113) | .501 (.722) | .578 (.318) |
| | 2 | Inferior Parietal Lobule | Right | .359 (.871) | .739 (.113) | .819 (.203) |
| | 3 | Declive | Right | .963 (.002) | .760 (.280) | .896 (.017) |
| L Insula | 4 | Thalamus | Left | .929 (.008) | .241 (1.44) | .888 (.120) |
| | 5 | Inferior Parietal Lobule | Right | .438 (.621) | .890 (.020) | .941 (.061) |
| R Insula | 6 | Thalamus | Left | .671 (.185) | .754 (.100) | .622 (.490) |
| | 7 | Superior Temporal Gyrus | Left | .459 (.563) | .191 (1.80) | .396 (.974) |
| | 8 | Inferior Temporal Gyrus | Left | .312 (1.06) | .082 (3.26) | .661 (.424) |
| L Cerebellar Tonsil | 9 | Parahippocampal Gyrus | Left | .248 (1.40) | .086 (2.85) | .285 (1.19) |
| R Claustrum | 10 | Cingulate Gyrus | Left | .151 (2.19) | .362 (.860) | .624 (.485) |
| | 11 | Inferior Parietal Lobule | Right | ○- | .008 (8.22) | .596 (.533) |
| | 12 | Inferior Frontal Gyrus | Left | .633 (.233) | .162 (2.04) | .550 (.368) |
| | 13 | Superior Temporal Gyrus | Left | ○- | .030 (5.26) | .793 (.235) |
| | 14 | Inferior Parietal Lobule | Left | ○- | .046 (4.37) | .733 (.316) |
| L Cingulate Gyrus | 15 | Insula | Right | .542 (.381) | .671 (4.08) | .973 (.001) |
| L Superior Frontal Gyrus | 16 | Medial Frontal Gyrus | Left | ○- | .022 (6.11) | .367 (1.07) |
| | 17 | Middle Temporal Gyrus | Left | .88 (3.28) | .525 (.420) | .744 (.301) |
| | 18 | Lentiform Nucleus | Left | .130 (2.47) | .227 (1.54) | .518 (.683) |
| R Inferior Frontal Gyrus | 19 | Inferior Frontal Gyrus | Left | .485 (.502) | .364 (.857) | .418 (.921) |

The effect of head motion (maximum framewise displacement [FD]) on functional connectivity was statistically assessed by using a linear mixed-effects model, in which a within-participants fixed effect of FD and interactions of FD × PHASE and FD × GROUP were included, in addition to the fixed effects of PHASE, GROUP, and PHASE × GROUP and a random effect of each individual participant. If the head movement-related interactions (i.e., FD × PHASE and FD × GROUP) were statistically significant, the statistics for FD were omitted because it was difficult to determine the effect.

### Distance matrices of the 12 brain regions showing recovery-selective increases in connectedness.

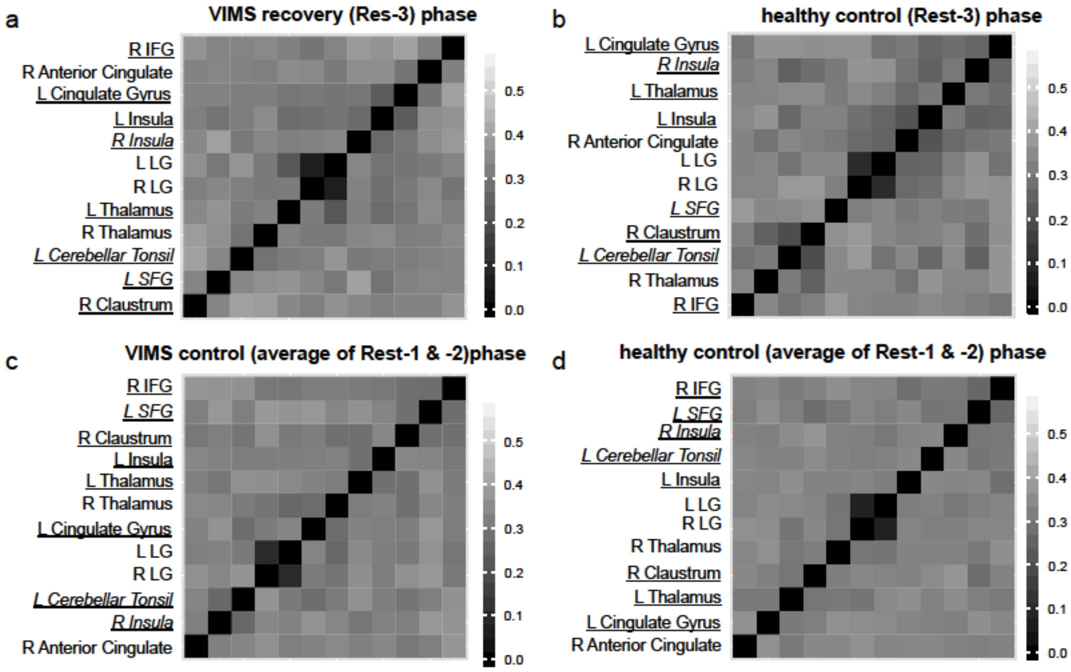


**Supplementary Figure 2. Distance matrices of the 12 brain regions showing recovery-selective increases in connectedness.**

(a) The distance matrix of the VIMS group for the recovery phase (Rest-3), whose elements indicate 1 minus absolute partial correlation between each pair of the 12 regions. Brain regions showing recovery-selective increases in the ROI-based functional connectivity and correlations with SSQ are underlined and in italics, respectively. (b) The same as (a) but for the healthy group. (c) The same as (a) but for the control phase (the average of Rest-1 and Rest-2). (d) The same as (c) but for the healthy group. Abbreviations: L, left; R, right; LG, lingual gyrus; SFG, superior frontal gyrus; IFG, inferior frontal gyrus.

### Extraction of independent brain networks with dictionary learning

To complement our main results, we performed a separate network analysis that extracted statistically independent resting networks related to the recovery from VIMS. A dictionary

learning technique (Mensch et al. 2016) was applied for the fMRI data of the individuals in the VIMS group, using the *nilearn* module of Python (https://nilearn.github.io/). First, after preprocessing of the fMRI data as described in the main text, the data during the recovery phase were subjected to dictionary learning (*DictLearn* and *RegionExtractor* methods with the number of components = 15 and minimum region size = 100 voxels), revealing 22 sparse regions of brain activities. Second, for each pair of the 22 brain regions, cross-correlation coefficients of fMRI time-series were computed, leading to a 22 × 22 matrix. Finally, this matrix was subjected to a connectome analysis using the *plot_connectome* function (edge threshold = 90%) in *nilearn*.
The result is shown in Supplementary Figure 3. Multiple brain regions critical for VIMS dynamics were found. The connectome map (Supplementary Figure 3b) shows visual processing regions, including the middle temporal gyrus and lingual gyrus, which were also found in the connectedness/functional connectivity approach used in the main text. These visual regions have been reported to have connectivity changes in the evolutional phase of VIMS (Miyazaki *et al.* 2015; Toschi *et al.* 2017). In addition, the cingulate region, which has been indicated to play key roles in the interoceptive process, such as self-awareness to one's own unpleasant bodily state, during/after VIMS, was detected in this analysis as well as that of the main text (Tables 2–5). Interestingly, the parahippocampal gyrus was again extracted in this analysis. The parahippocampal region is a core area for spatial memory processing (Epstein *et al.* 1998; Epstein, 2008), and memory processing has been suggested to be involved in an adaptation to motion sickness including VIMS (Reason & Brand 1975; Reason 1978; Oman 1990). This result thus suggests that such an adaptive process might underlie the recovery from VIMS. Altogether, this analysis corroborates our results reported in the main text.

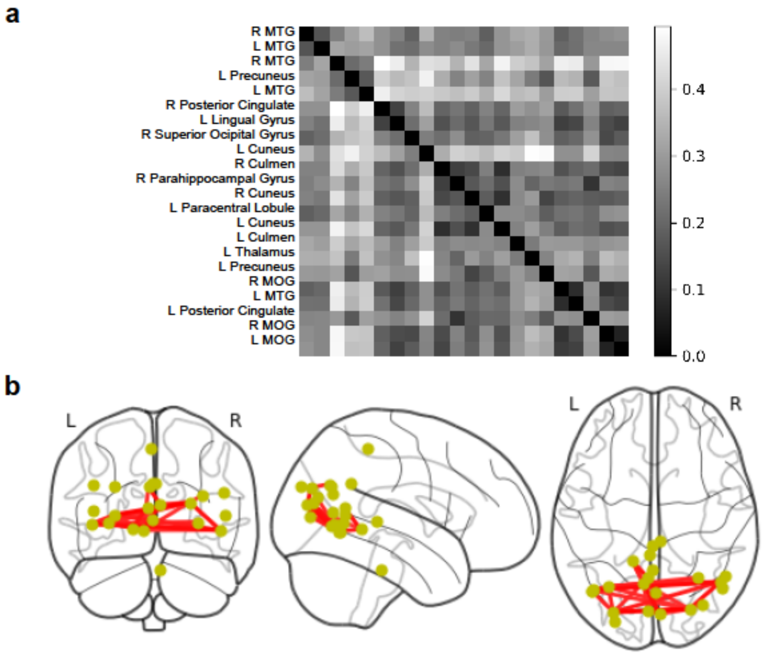


**Supplementary Figure 3. Statistically independent brain networks related to recovery from visually induced motion sickness (VIMS)**

(a) The within-participant averaged distance matrix for the recovery phase of the VIMS group. Each element represents a cross-correlation coefficient between each pair of the 22 regions determined by a dictionary learning technique. This analysis derived 22 regions of interest (ROIs). (b) The top 10% connections derived from the distance matrix. The connections are overlaid on a glass brain. Yellow circles and red lines denote the derived regions and connections, respectively. The connections are as follows: R Posterior Cingulate - L Lingual Gyrus, L Lingual Gyrus - R MOG, L Lingual Gyrus - R MOG, L Lingual Gyrus - L MTG, L Lingual Gyrus - R MOG, L Lingual Gyrus - L MOG, R Superior Ocipital Gyrus - R MOG, R Culmen - R Parahippocampal Gyrus, R Culmen - R Cuneus, R Culmen - L Cuneus, R Culmen - R MOG, R Culmen - L MOG, R Parahippocampal Gyrus - L Cuneus, R Parahippocampal Gyrus - L Posterior Cingulate, R Cuneus - L Cuneus, R Cuneus - L Precuneus, L Cuneus - R MOG, R MOG - L MTG, L MOG - R MOG, L MOG - R MOG, L MTG - R MOG, L MTG - L MOG, and R MOG - R MOG. Abbreviations: L, left; R, right; MTG, middle temporal gyrus; MOG, middle occipital gyrus.
